# Programmable genetic control of tumor-colonizing *Bifidobacterium longum* for intratumoral therapeutic delivery and biocontainment

**DOI:** 10.64898/2026.08.12.744520

**Authors:** Jaehyun Lee, Joshua Glazier, Ralph R. Weichselbaum, Mark Mimee

**Affiliations:** Department of Microbiology, University of Chicago, Chicago, IL 60637, USA; Pritzker School of Molecular Engineering, University of Chicago, Chicago, IL 60637, USA; Department of Radiation and Cellular Oncology, University of Chicago, Chicago, 60637, USA; The Ludwig Center for Metastasis Research, University of Chicago, Chicago, 60637, USA; Duchoissois Family Institute, University of Chicago, Chicago, IL 60637, USA

## Abstract

Engineered bacteria offer a distinct modality for cancer therapy by exploiting the ability of certain species to colonize tumors and deliver therapeutic payloads. Improving their efficacy and safety requires control over bacterial activity after tumor colonization, yet few microbial chassis permit it. *Bifidobacterium longum*, a probiotic with intrinsic tumor-targeting and antitumor activity, is a promising chassis but lacks such control. Here, we develop a genetic control system that regulates *B. longum* activity within tumors, from gene expression to bacterial abundance. A human-isolate-derived replicon supports plasmid maintenance without antibiotic selection, and promoter and ribosome-binding-site libraries provide ∼150-fold and ∼48-fold expression ranges, respectively. Signal peptides enable secretion of structurally diverse therapeutic payloads and *B. longum* secreting CCL21 or an anti-PD-L1 nanobody reduces tumor growth relative to PBS controls. Anhydrotetracycline delivered in drinking water induces transgene expression in tumor-resident bacteria and reduces intratumoral bacterial load through CRISPRi targeting essential genes. Together, these results establish a tumor-homing probiotic as an externally controllable therapeutic chassis.

## Introduction

Bacterial cancer therapy exploits the natural propensity of certain bacteria to colonize within tumors and perform therapeutic functions at the site of disease^1–4^. Synthetic biology has extended this promise, turning bacteria into targeted delivery platforms that produce therapeutic molecules such as immunomodulatory cytokines^5^ and chemokines^6,7^ and checkpoint-blocking nanobodies^8,9^. Yet effective and safe therapy requires not only localized delivery, but also control over gene expression, payload release, and bacterial growth in situ. Intratumoral control has been developed most extensively in highly genetically tractable chassis, such as *Escherichia coli* and *Salmonella* Typhimurium, for which rich collections of genetic parts have been characterized. Chemical^10,11^-and focused-ultrasound^12^-based induction systems regulate payload expression on demand within tumors. Quorum-lysis circuits couple population density to payload release^8^, while nutrient auxotrophies and hypoxia-sensing circuits confine proliferation of engineered microbes to the tumor^11,13^. Together, these strategies demonstrate that both bacterial gene expression and growth can be genetically regulated *in vivo*, offering important control elements to improve the safety and efficacy of engineered bacterial cancer therapy.

Beyond genetic tractability, an ideal chassis for cancer therapy would be an obligately anaerobic, favoring growth in hypoxic tumors over oxygenated healthy tissues, and non-pathogenic in origin^2,3^. Together, these properties would intrinsically confer tumor selectivity and safety, reducing reliance on extensive attenuation or containment engineering. *Bifidobacterium* spp. offer a compelling alternative to commonly used chassis, including *E. coli, S.* Typhimurium, and *Listeria monocytogenes*^14–17^. As obligate anaerobes and beneficial gut symbionts with a long probiotic history^14^, they exhibit high tumor selectivity with a favorable safety profile that obviates the attenuation required for pathogenic microorganisms^18–20^. *Bifidobacterium* can also support antitumor immunity within the tumor microenvironment (TME). For example, they can activate dendritic cells to enhance responses to CD47 checkpoint blockade^21^. However, their development into a programmable therapeutic platform remains constrained by a lack of versatile genetic engineering tools to enable robust intratumoral control of payload delivery and bacterial growth.

For *Bifidobacterium* to function as a programmable therapeutic chassis, genetic control must extend beyond constitutive transgene expression^22,23^. Therapeutic deployment requires stable maintenance of engineered programs, quantitative tuning of expression levels, localization of therapeutic proteins outside the bacterial cytoplasm, inducible regulation by an externally administered signal, and the ability to modulate bacterial growth or persistence after tumor colonization. Although heterologous inducible systems have been introduced into *Bifidobacterium*, their performance can be constrained by host-specific regulatory compatibility and metabolic requirements^24^. For example, the nisin-controlled expression system widely used in lactic acid bacteria shows substantial basal activity and limited induction in *B. longum* NCC2705^24^, and the *E. coli*-derived araC–P_BAD_ system is restricted to strains capable of arabinose uptake and utilization^25,26^. Critically, these systems have been characterized almost exclusively in culture, leaving it unclear whether external control persists after tumor colonization. For growth control, CRISPR interference (CRISPRi) has achieved growth inhibition in *B. breve*, but only by repressing genes required to metabolize the supplied carbohydrate, a dependency not controllable *in vivo*^27^.

Here, we establish external control over gene expression and growth in *B. longum* after tumor colonization. To support this control, we first established stable plasmid maintenance and calibrated gene expression by identifying a native replicon and constructing promoter and ribosome-binding-site (RBS) libraries. We then directed heterologous proteins to the extracellular milieu or cell surface using natural and synthetic signal peptides and cell-wall-anchoring motifs. Engineered *B. longum* secreted diverse therapeutic payloads, including cytokines, chemokines and immunomodulatory nanobodies, and strains secreting CCL21 or an anti-PD-L1 nanobody retained antitumor activity relative to PBS-treated controls *in vivo*. Finally, we achieved inducible control of gene expression and growth in tumor-resident bacteria by coupling heterologous expression and CRISPRi against essential genes to an anhydrotetracycline (aTc)-inducible system. Together, these capabilities support the use of *B. longum* as a programmable therapeutic chassis whose gene expression and growth remain controllable after tumor colonization.

## Results

### A native replicon supports stable genetic programming of *B. longum*

*Bifidobacterium* spp. are a promising chassis for live bacterial therapeutics, particularly for cancer therapy^28^. However, their genetic engineering has been limited by the lack of host-compatible replicons that support stable maintenance and expression of engineered constructs^29^. Reasoning that a native replicon would confer robust plasmid retention^30^, we surveyed *Bifidobacterium* plasmid sequences in NCBI and compared them with genomes from the symbiotic bacterial strain bank (Supplementary Fig. 1a, b). We identified a locus in *B. longum* MSK 17.29 sharing 99.5% and 90.5% sequence identity with the native plasmids pMG1 and pTB6, respectively, and isolated its origin of replication and cognate rep gene to establish a core replicon for *Bifidobacterium* (Supplementary Fig. 1c).

Using this replicon, we built a minimal expression vector carrying a chloramphenicol-resistance gene (*cat*) for selection and a NanoLuc luciferase reporter for luminescence-based quantification of heterologous gene expression (Fig. 1a). We generated two architectures. Shuttle vectors carried an additional *E. coli* origin for amplification in *E. coli* before transfer, whereas direct vectors omitted it and are transformed into *B. longum* immediately after assembly, shortening the overall workflow. Within each architecture, two variants differed in the promoter used to drive *cat* expression (P_tuf-Bb_ or P_hup_). Because P_hup_ is silent in *E. coli*, the P_hup_ shuttle vector carried a dedicated *E. coli* cassette (P_kanR_-driven kanamycin resistance). The resulting four vectors were introduced into *B. longum* ATCC 15697.

**Fig. 1.**
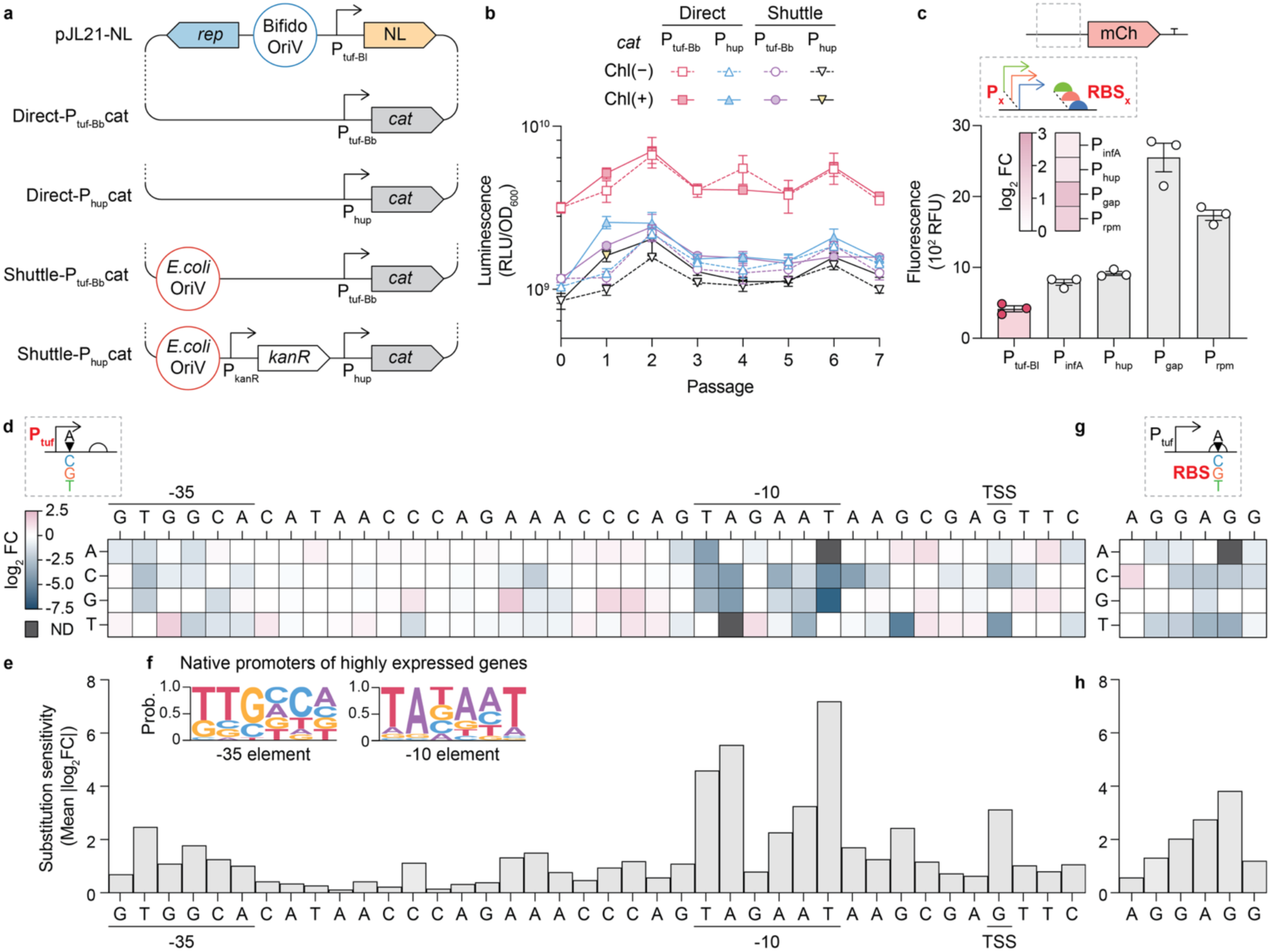
Plasmid constructs and synthetic promoter/RBS libraries for tunable expression in *B. longum*. (a) Schematic of the four plasmid constructs: two direct-expression vectors (P*_tuf_*_-Bb_ or P*_hup_* driving *cat*) and two corresponding *E. coli*–*B. longum* shuttle vectors. (b) Plasmid retention assay. Luminescence per OD_600_ of cultures over serial passages in the presence or absence of 5 µg/ml chloramphenicol (n = 4). (c) mCherry (mCh) expression driven by native *B. longum* promoters from five housekeeping genes (*tuf, rpmB, gap, hup and infA*) placed upstream of an mCherry reporter (n = 3). Bar graph, fluorescence; heatmap, mean log_2_ fold change (log_2_FC) in fluorescence relative to the *tuf* promoter. Promoter sequences and transcription start sites (TSS) are provided in Supplementary Table 2. (d) Effect of single-nucleotide substitutions across the *tuf* promoter on expression, shown as a heatmap of mean log2 fold change relative to wild-type. Columns, promoter position labeled by the wild-type nucleotide; rows, substituted nucleotide, with wild-type positions shown in white (n = 3–4). ND, not detected (fluorescence was at or below the level of the non-fluorescent background control). (e) Substitution sensitivity at each position of the *tuf* promoter, calculated as the mean of the absolute per-substitution log_2_FC values across all single-nucleotide substitutions at that position (n = 3–4). (f) Sequence logos of the −35 and −10 promoter elements, derived from 19 and 24 promoters, respectively, that are associated with highly expressed genes. (g) Effect of single-nucleotide substitutions across the *tuf* ribosome-binding site (RBS) on expression, shown as a heatmap as in (d) (n = 3). (h) Substitution sensitivity at each position of the *tuf* RBS, calculated as in (e) (n = 3). Data in (b) and (c) are mean ± s.e.m.

Because stable plasmid retention without antibiotics is crucial for *in vivo* use, we serially passaged strains carrying each of the four vectors with or without antibiotic selection (Fig. 1b). All strains maintained stable luminescence over seven passages (∼77 generations) regardless of selection. Comparable colony-forming unit (CFU) counts on selective and non-selective plates after the final passage of unselected cultures confirmed stable replicon retention without selective pressure (Supplementary Fig. 1d).

Vector architecture strongly influenced expression (Supplementary Fig. 1e, f). Direct vectors produced higher luciferase activity and tended to have higher plasmid copy numbers than their matched shuttle counterparts. Notably, the P_hup_ shuttle vector, the conventional dual-selection architecture for non-model bacteria^31^, showed the lowest luciferase activity and copy number among the four.

Across *Bifidobacterium* spp., the plasmid was established and generated detectable luminescence in four of six *B. longum* strains and one of two *B. pseudocatenulatum* strains, including human-derived isolates, but not in tested strains of the other species (Supplementary Fig. 1g, h).

Together, these results establish a stable, *Bifidobacterium*-derived plasmid system that supports reliable maintenance and expression of heterologous constructs for payload delivery and biocontrol.

### Synthetic promoter and RBS libraries enable quantitative expression tuning in *B. longum*

Having established stable plasmid maintenance in *B. longum*, we next sought to tune gene expression from these plasmids. Quantitative control is important for therapeutic applications because engineered functions must achieve sufficient activity without imposing excessive burden on the bacterial host^32^. As a starting point, we cloned the putative promoter regions (121–155 bp upstream of the start codon) of five housekeeping genes (*tuf*, *rpmB*, *gap*, *hup*, and *infA*) in front of an mCherry reporter (Fig. 1c). These native promoters drove distinct levels of constitutive expression but spanned only a narrow range (∼6-fold). To expand this range, we generated a library of 120 P*_tuf_* variants containing single-nucleotide substitutions across the −35 and −10 elements, the transcription start site (TSS), and intervening regions. These spanned an ∼150-fold range and achieved expression up to ∼4.6-fold higher than native P*_tuf_*, extending the tunable range beyond the native set (Fig. 1d, Supplementary Fig. 2a). Substitution sensitivity varied across the promoter, being highest in the −10 element, lower in the −35 element and TSS-proximal region, and relatively minor across the spacer (Fig. 1e). To assess the biological relevance of the substitution landscape, we analyzed the highly expressed native promoters of *B. longum* ATCC15697, recovering the canonical *Bifidobacterium* −35 (TTGNNN) and −10 (TANNNT) motifs^33^ (Fig. 1f). The −10 element, the most substitution-sensitive position, was also more conserved in native promoters, while the −35 element, with smaller substitution effects, showed weaker conservation, supporting the biological relevance of the inferred landscape.

To extend tunable control to the translational level, we applied the same single-nucleotide substitution strategy to the RBS*_tuf_* (Fig. 1h, Supplementary Fig. 2b). The RBS library spanned an ∼48-fold range, and the largest changes arose within the central GGAG segment (Fig. 2i). Expression negatively correlated with predicted RBS-16S rRNA hybridization energy (Spearman’s *r* = −0.72, *p* = 0.0004), consistent with stronger complementarity generally enhancing translation^34^ (Supplementary Fig. 2c, d). However, RBS*_tuf_*_-A1C_ (CGGAGG) outperformed native AGGAGG despite less favorable predicted hybridization, suggesting that optimal rather than maximal complementarity drives better translation, consistent with prior observations in *Bifidobacterium*^35^. Together, these libraries enabled graded transcriptional and translational expression control in *B. longum*.

**Fig. 2.**
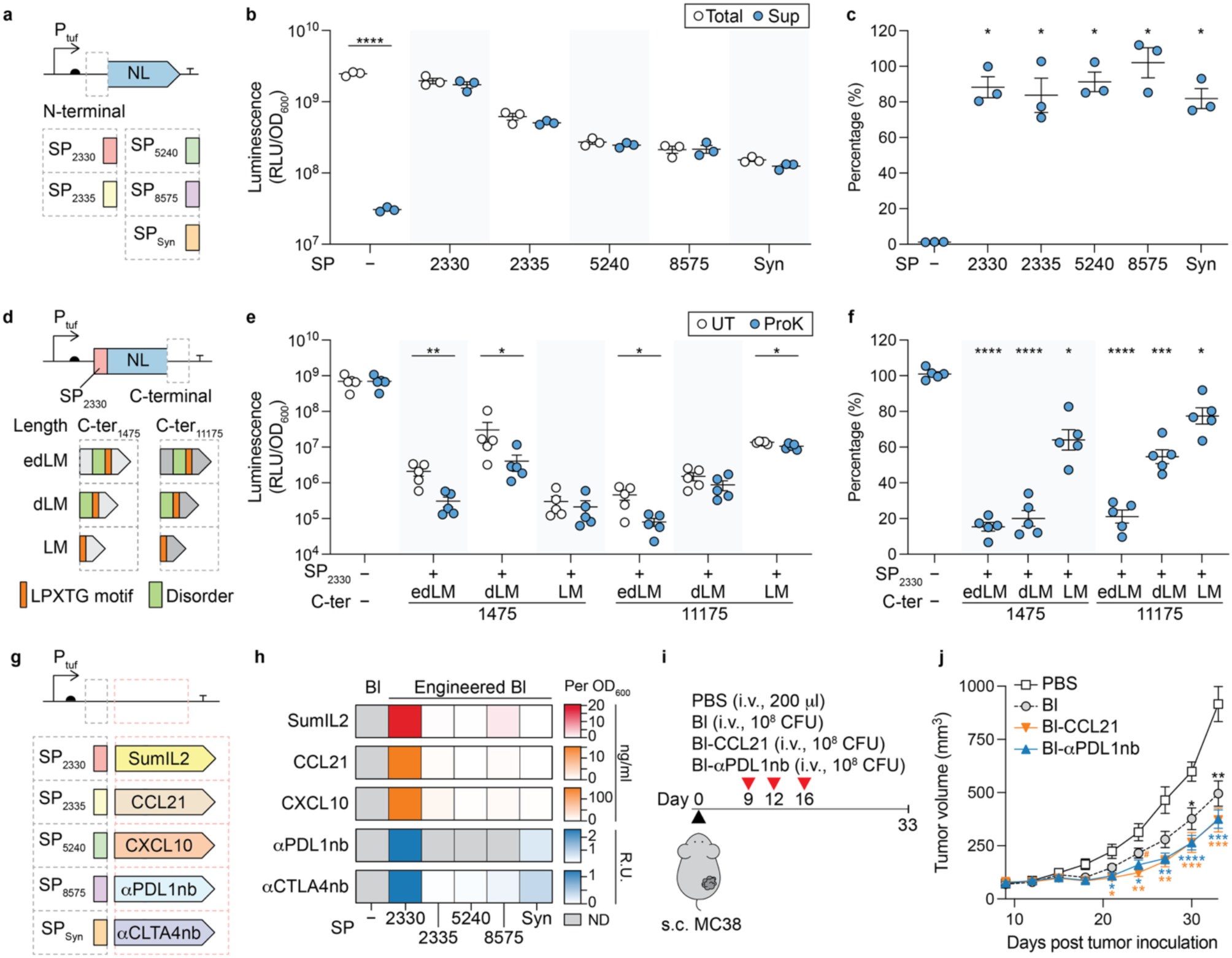
Secretion and surface-display modules for controlled localization and secretion of diverse therapeutic proteins in *B. longum*. (a) Schematic of N-terminal signal peptides (SPs) fused upstream of NanoLuc luciferase (NL), driven by the P*_tuf_* promoter. Four native SPs (SP_2330_, SP_2335_, SP_5240_, and SP_8575_) and one synthetic SP (SP_syn_) were tested, along with a control strain lacking a signal peptide. SP_syn_ was derived from a consensus of 27 natural SPs (Supplementary Fig. 3). (b,c) Luminescence per OD_600_ in whole culture (Total) and supernatant (Sup) for the strains tested (b), and the percentage of secreted luminescence, calculated as (Sup/Total) × 100 (c) (n = 3). (b) Welch’s t-test on log-transformed data. (c) Brown–Forsythe and Welch ANOVA followed by Dunnett’s T3 multiple-comparison test against the control strain. (d) Schematic of C-terminal cell-wall–anchoring fragments fused downstream of NL with the SP_2330_ signal peptide, for surface display. Fragments were derived from two LPXTG-containing cell-wall proteins, yielding the C-ter_1475_ and C-ter_11175_ series. Within each series, three fragments of increasing length were tested: a minimal LPXTG motif-containing region (LM), a version additionally including the predicted disordered region (dLM), and a longer upstream extension (edLM). (e,f) Luminescence per OD_600_ in proteinase K–treated (ProK) and untreated (UT) strains harboring SP_2330_ and a C-terminal LPXTG motif, and a control strain lacking both (e), and the percentage of luminescence remaining after proteinase K treatment, calculated as (ProK/UT) × 100 (f) (n = 5). (e) Welch’s t-test on log-transformed data. (f) Brown–Forsythe and Welch ANOVA followed by Dunnett’s T3 multiple-comparison test against the control strain. (g) Schematic of the signal peptide (SP) and therapeutic protein combinations tested: Supermutant IL-2 (SumIL2), CXCL10, CCL21, an anti–PD-L1 nanobody (αPDL1nb) and anti–CTLA-4 nanobody (αCTLA4nb). (h) Secretion of therapeutic proteins into culture supernatants, measured by ELISA and shown as a heatmap of mean values (n = 3). ND, not detected. (i) Intravenous treatment of mice bearing established subcutaneous MC38 tumors with PBS, wild-type *B. longum* (Bl; 10^8^ CFU), Bl-CCL21 (10^8^ CFU) or Bl-αPDL1nb (10^8^ CFU) on days 9, 12, and 16. (j) Mean tumor volume over time (n = 11 per group, pooled from two independent experiments). Two-way repeated-measures ANOVA followed by Tukey’s multiple-comparison test. Asterisks, each group versus PBS. #, P < 0.05 for Bl versus Bl.CCL21. Data in (b), (c), (e), (f) and (j) are mean ± s.e.m. *P < 0.05, **P < 0.01, ***P < 0.001, ****P < 0.0001.

### Secretion and surface-display modules for controlled localization of heterologous proteins

Beyond expression level, therapeutic activity also depends on protein localization, particularly when the intended target is located outside the bacterial cell^36^. We therefore established modules that direct heterologous proteins to the cell exterior, either through secretion or surface display. For secretion, we used SignalP6^37^ to screen the *B. longum* proteome for native signal peptides predicted to direct proteins through the Sec pathway. We selected four high-scoring native peptides (>0.99) for direct use and designed a synthetic peptide from the consensus of 27 high-scoring native candidates (>0.99) (Fig. 2a, Supplementary Fig. 3). Each signal peptide was fused to the N-terminus of NanoLuc, and secretion was quantified by comparing luciferase activity in cell-free supernatants to that in whole cultures (Fig. 2b, c). Although total luciferase activity varied across signal peptides, all five peptides supported efficient extracellular accumulation, with an average of more than 80% of total luminescence detected in the supernatant. In contrast, luciferase lacking a signal peptide yielded only ∼1% of total luminescence in the supernatant. Thus, both native and synthetic signal peptides support efficient protein secretion in *B. longum*.

For surface display, we leveraged LPXTG cell-wall anchoring sequences, which mediate sortase-dependent anchoring in Gram-positive bacteria^38^. We identified two proteins (BLON_RS01475, BLON_RS11175) containing LPXTG motifs from UniProt annotation^39^, and, for each, designed three C-terminal fragments of increasing length: a minimal motif-containing region (LM), an extension including the predicted disordered region (dLM), and a longer upstream extension (edLM) (Fig. 2d). Each fragment was fused to the C-terminus of NanoLuc, carrying the SP_2330_ signal peptide to promote export from the cytoplasm. We first estimated the secreted fraction, reasoning that efficiently surface-displayed NanoLuc should remain cell-associated rather than accumulate in the supernatant (Supplementary Fig. 4). The edLM fusions were largely secreted (∼90% in the supernatant), whereas LM and dLM fusions remained more cell-associated, suggesting that the additional upstream extension did not improve cell association. However, cell-associated signals cannot distinguish surface-displayed from intracellularly retained protein. We therefore directly assessed surface exposure directly by treating washed cells with proteinase K (Fig. 2e, f). Luminescence from the construct lacking both a signal peptide and an LPXTG motif was unaffected by proteinase K treatment, which confirmed that the intracellular luciferase was inaccessible to the protease under these conditions. Several constructs (C-ter_1475-LM_, C-ter_11175-dLM_ and C-ter_11175-LM_) retained >50% of their luminescence after treatment, indicating that a substantial fraction of the reporter remained intracellular. By contrast, C-ter_1475-edLM_, C-ter_1475-dLM_ and C-ter_11175-edLM_ showed substantial luminescence loss, supporting their surface display. Among these, C-ter_1475-dLM_ showed the strongest surface-display phenotype, with low supernatant luminescence and high protease sensitivity. Nonetheless, residual luminescence persisted across constructs after treatment, which points to room for further optimization. Together, these two strategies, signal peptide–mediated secretion and LPXTG-based surface display, supported external localization of heterologous proteins in *B. longum*.

### Engineered *B. longum* secretes structurally diverse therapeutic proteins while retaining antitumor activity ***in vivo***

We next examined whether signal peptide–mediated secretion could be extended to structurally diverse therapeutic proteins and how secretion depended on signal peptide–payload pairing (Fig. 2g). We first fused each of the five validated signal peptides to Supermutant IL-2 (SumIL2), an engineered immunostimulatory IL-2 variant^40^ previously shown to enhance antitumor efficacy when secreted by *B. longum* in both subcutaneous and orthotopic pancreatic tumor models^17^. Extracellular SumIL2 levels varied substantially across signal peptides, with SP_2330_ supporting the highest secretion (Fig. 2h).

We extended this to four additional antitumor payloads, each previously expressed in other engineered bacterial chassis: chemokines that recruit immune cells into tumors (CCL21^7^, CXCL10^41^) and immune checkpoint– targeting nanobodies (anti–PD-L1nb^8^, anti–CTLA-4nb^8^) (Fig. 2h). As with SumIL2, secretion levels varied across combinations. SP_2330_ again supported the highest secretion, extending its advantage to all four additional payloads tested, whereas the relative performance of the other signal peptides was payload dependent. For example, SP_Syn_ was the second-most effective signal peptide for anti-PD-L1nb and anti-CTLA-4nb secretion but performed poorly for CCL21 and CXCL10 secretion. At the extreme, anti-PD-L1nb was not detected when combined with SP_2335_, SP_5240_ and SP_8575_, indicating that signal peptide selection can abolish detectable payload secretion.

We then evaluated whether engineered strains retain antitumor activity *in vivo*, using two strains secreting payloads from distinct classes: CCL21 (Bl-CCL21) and an anti–PD-L1 nanobody (Bl-αPDL1nb). Mice bearing established subcutaneous MC38 tumors received intravenous PBS, wild-type *B. longum*, Bl-CCL21, or Bl-αPDL1nb every 3–4 days (Fig. 2i). Both engineered strains significantly reduced tumor growth relative to the PBS control from day 21 (Fig. 2j). Consistent with its intrinsic antitumor properties^17^, wild-type *B. longum* also showed significant antitumor activity relative to the PBS control, but only from day 33. Tumor volumes were significantly lower in Bl-CCL21–treated mice than in wild-type–treated mice at day 24, although this difference was not sustained at later time points.

Together, these signal peptides supported pairing-dependent secretion of structurally diverse therapeutic proteins from *B. longum*, while payload-secreting strains retained antitumor activity *in vivo*.

### A tunable inducible system controls *B. longum* gene expression in tumors and in the gut

Beyond constitutive expression, inducible control would make *B. longum* a more versatile therapeutic chassis by allowing engineered functions to be activated only when needed. We therefore built an inducible system using the TetR/tetO module, chosen because its inducer aTc enters cells by diffusion and acts directly on TetR without requiring metabolic activation^42^, thereby rendering induction independent of strain-specific metabolic capacity. We constitutively expressed *tetR* from P_infA_ and evaluated three configurations for tetO placement within the P_tuf_: upstream of the −35 element (distal; P*_tuf_*_(tetO-d)_), downstream of the TSS (proximal; P*_tuf_*_(tetO-p)_), or at both positions (P*_tuf_*_(tetO-pd)_) (Fig. 3a). We excluded placement within the −35/−10 spacer because insertion of the 19bp operator would disrupt core promoter elements, whose substitution markedly reduced activity in our mutagenesis screen (Fig. 1d, e). Each variant was fused to NanoLuc and induced with aTc. Without TetR, aTc did not affect luciferase activity from any variant, confirming that tetO alone does not confer inducibility (Fig. 3b, c). With TetR, repression depended on tetO position. Native P*_tuf_* and the distal variant showed little induction, whereas the proximal variant was repressed in the absence of aTc and significantly induced upon its addition. The dual-site variant was only modestly induced, possibly because the distal operator diverted TetR away from the proximal operator. Thus, a tetO site downstream of the TSS is required for effective TetR-mediated repression. Dose-response analysis of the proximal variant defined a functional range of 7.8–500 ng/ml aTc (Fig. 3d). We then broadened the dynamic range, using the genetic elements characterized above (Fig. 1d, g and 2e). Replacing P*_infA_* with the stronger P*_gap_* promoter to drive *tetR* expression raised the induction ratio from ∼9 to ∼100-fold (Fig. 2f). Weakening reporter translation with RBS variants (G3C < G6T < native AGGAGG) shifted the dose-response curve downward, consistent with their characterized strengths (Fig. 1d, g). Combined, these two modifications lowered the basal expression level by up to ∼1000-fold relative to the original system. Together, these results establish a tunable aTc-inducible expression system for *B. longum*.

**Fig. 3.**
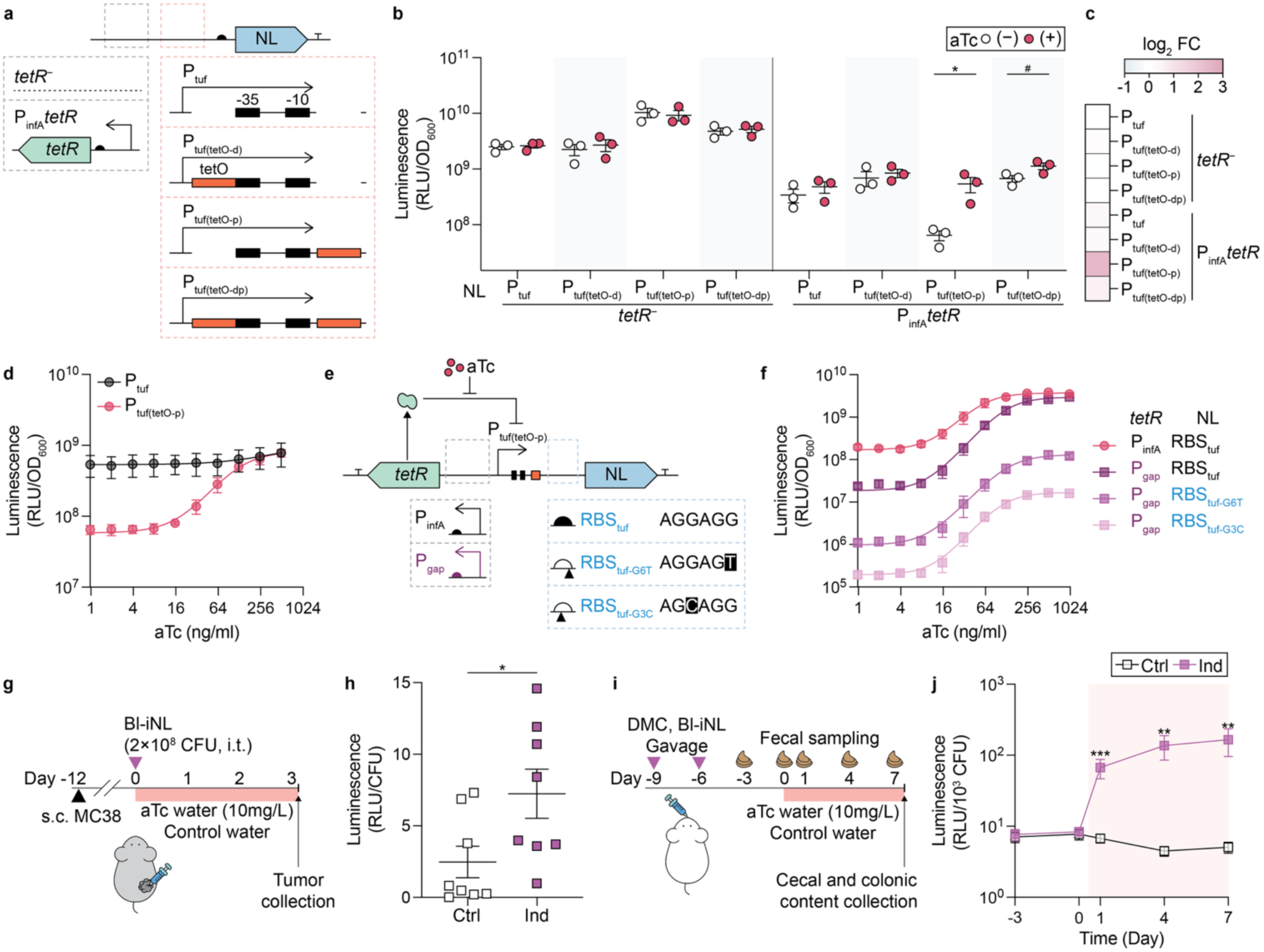
A tunable, aTc-inducible expression system controls *B. longum* gene expression in tumors and in the gut. (a) Design of the inducible constructs, comprising P*_tuf_*_-tetO_ promoter variants with or without *tetR* expressed from P*_infA_*. (b,c) Luminescence per OD_600_ in culture in the presence or absence of 250 ng/ml aTc, with or without co-expression of tetR (n = 3). (b) Welch’s t-test on log-transformed data. (c) Heatmap of the mean log_2_FC in luminescence (aTc+/aTc−). #P = 0.069 (d) aTc dose–response of luminescence per OD_600_ for the P*_tuf_* and P*_tuf_*_-tetO-p_ promoters, fitted with a four-parameter logistic model (n = 3). (e) Tuning of the inducible system by replacing P_infA_ with P_gap_ for *tetR* expression and varying the RBS to shift the dynamic range. (f) aTc dose–response curves of luminescence per OD_600_ for the P*_gap_*-_tetR_ inducible variants with different RBSs, fitted with a four-parameter logistic model (n = 3). (g) Intratumoral injection of Bl-iNL (P*_gap_*-*tetR*, P*_tuf_*_(tetO-p)-RBS*tuf*-G6T_-NL, 2 × 10^8^ CFU) into established subcutaneous MC38 tumors, followed by aTc-containing (10 mg/L, Ind) or control (Ctrl) drinking water. Tumors were collected 3 days after injection. (h) Luminescence per CFU in tumors (n = 8, pooled from two independent experiments). Welch’s t-test. (i) Gavage of germ-free mice with Bl-iNL (3 × 10^9^ CFU) and a defined microbial community (DMC) on days −9 and −6, followed by aTc-containing (10 mg/L, Ind) or control (Ctrl) drinking water from day 0 to day 7. Fecal samples were collected longitudinally (days −3 to 7), and cecal and colonic contents at the endpoint. (j) Luminescence per CFU in feces over time (n = 5 for Ctrl and n = 7 for Ind, pooled from two independent experiments). Ctrl and Ind were compared at each time point by two-way repeated-measures ANOVA on log-transformed data followed by Šídák’s multiple-comparison test. (g-j) See Supplementary Fig. 5 for bacterial burden and luciferase activity. Data in (b), (d), (f), (h) and (j) are mean ± s.e.m. *P < 0.05, **P < 0.01, ***P < 0.001, ****P < 0.0001.

To test whether this system remains externally controllable within tumors, we injected Bl-iNL (P_gap_-tetR, P_tuf(tetO-p)_-RBS_tuf-G6T_-NL) intratumorally into established subcutaneous MC38 tumors and simultaneously provided aTc-containing or control drinking water (Fig. 3g). Three days after bacterial injection, bacterial loads were comparable between aTc-treated and control groups, indicating that aTc did not detectably affect bacterial abundance within tumors at this time point (Supplementary Fig. 5a). Despite these comparable burdens, luminescence per CFU was significantly higher in the aTc-treated group (2.9-fold; Fig. 4h), demonstrating that an orally administered inducer can activate bacterial gene expression within tumors.

**Fig. 4.**
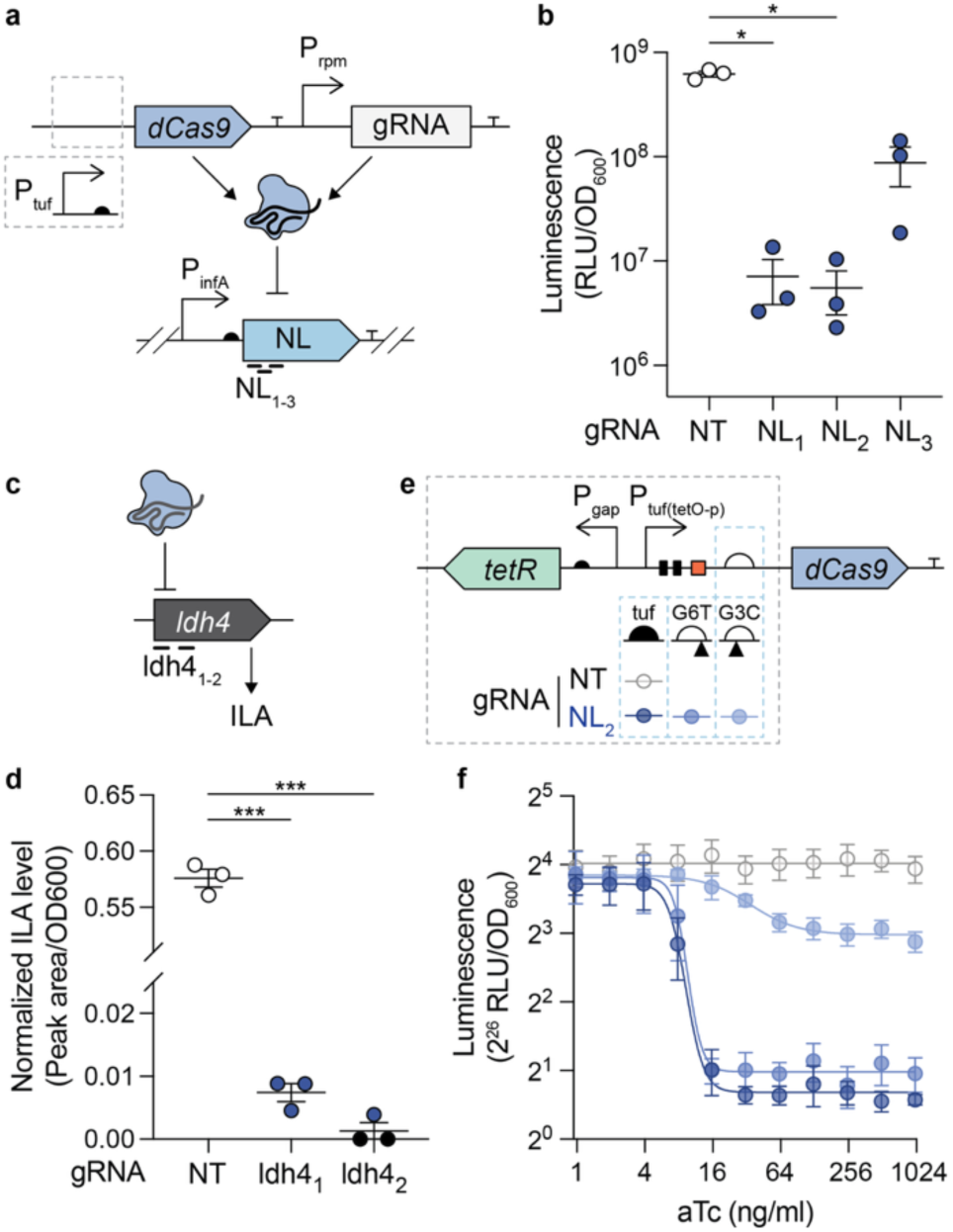
A tunable CRISPRi system for gene knockdown in *B. longum*. (a) Design of the CRISPRi system. dCas9 and the gRNA are constitutively expressed from P*_tuf_* and P*_rpmB_*, respectively. A NanoLuc luciferase (NL) reporter within the same construct serves as the repression target. (b) CRISPRi repression of the NL reporter by three gRNAs, relative to a non-targeting control (NT), measured as luminescence per OD_600_ (n = 3). Brown–Forsythe and Welch ANOVA on log-transformed data followed by Games–Howell’s multiple-comparison test. (c) CRISPRi targeting of *ldh4* (BLON_RS05510), which produces indole-3-lactic acid (ILA), with two gRNAs. (d) Resulting ILA levels (peak area per OD_600_), compared against NT (n = 3). Brown–Forsythe and Welch ANOVA followed by Dunnett’s T3 multiple-comparison test. Data points on the x-axis indicate samples in which ILA was not detected (peak area = 0). (e) Inducible CRISPRi system, in which dCas9 expression is controlled by aTc and tuned with different RBSs (native, G6T and G3C). A non-targeting (NT) gRNA with the native RBS serves as a control. (f) aTc dose–inhibition curves of luminescence per OD_600_ for the inducible CRISPRi variants with different dCas9 RBSs and a non-targeting (NT) control, all fitted with a four-parameter logistic model (n = 3). Data are mean ± s.e.m. *P < 0.05, **P < 0.01, ***P < 0.001, ****P < 0.0001.

To determine whether this control extended beyond tumors, we next evaluated the system in the gut, a native niche of *B. longum*^43^ and a physiologically distinct environment shaped by microbial and host interactions^44^. Germ-free mice were gavaged with Bl-iNL together with a defined microbial community (DMC)^45^ on days −9 and −6 and were subsequently provided aTc-containing or control drinking water from day 0 through day 7 (Fig. 3i). To quantify Bl-iNL abundance and luciferase activity, fecal samples were collected longitudinally from day −3 through day 7, and cecal and colonic contents were collected at the endpoint. Fecal bacterial abundance remained stable from day −3 through day 7 and did not differ between aTc-treated and control mice (Supplementary Fig. 5c). Before induction, fecal luminescence per CFU was comparable between groups on days −3 and 0 (Fig. 3j). Following aTc administration, luminescence per CFU increased sharply and was significantly higher than in controls at each post-induction time point (9.6-, 29.2-and 31.1-fold on days 1, 4 and 7, respectively; Fig. 3j). At the endpoint, cecal and colonic contents from aTc-treated mice likewise showed significantly higher luminescence per CFU than controls (∼17-fold in both compartments), while bacterial abundance remained comparable between groups (Supplementary Fig. 5e-j).

Together, these results establish that the TetR/tetO inducible system enables sustained external control of *B. longum* gene expression across distinct *in vivo* environments, the tumor and the gut, without measurably altering bacterial abundance.

### CRISPRi enables programmable control of heterologous and endogenous genes in *B. longum*

Although heterologous gene expression enables the installation of new therapeutic functions^3^, the outcome of bacterial cancer therapy is also shaped by the intrinsic biology of the bacterial chassis^1,2^, including its capacity for tumor colonization, metabolic activity and intrinsic antitumor activity — the last observed for wild-type *B. longum* (Fig. 2j). Regulating endogenous genes would therefore provide a means to investigate and manipulate native bacterial functions that influence both antitumor activity and the behavior of the chassis itself. Previous transposon mutagenesis studies in *B. breve* have enabled genome-wide analysis of native gene functions^46^, but their reliance on disruptive insertions limits functional analysis to mutants that can be recovered and does not allow specific genes to be selectively repressed^47^. CRISPRi, by contrast, enables target-specific repression without permanent genome disruption^48^.

We therefore established CRISPRi in *B. longum* by expressing nuclease-deactivated Cas9 (dCas9) and target-specific guide RNAs (gRNAs) from the constitutive promoters P*_tuf_* and P*_rpmB_*, respectively, on a single plasmid (Fig. 4a). We first targeted a NanoLuc reporter encoded on the same plasmid, and three independent gRNAs each mediated repression, with the most effective guide reducing luminescence ∼100-fold relative to a non-targeting (scrambled) gRNA control (Fig. 4b).

As an endogenous target, we selected *ldh4*, which encodes an aromatic lactate dehydrogenase involved in the production of indole-3-lactic acid (ILA)^49^. ILA is a tryptophan-derived aryl hydrocarbon receptor ligand with context-dependent immunomodulatory activity^50^ and may influence immune responses within the TME^51^. Two independent gRNAs targeting *ldh4* significantly reduced ILA production relative to a non-targeting control, demonstrating that CRISPRi can effectively silence an endogenous bacterial gene and modulate the production of a bacterially derived immunomodulatory metabolite (Fig. 4c, d).

Next, to enable external control of CRISPRi-mediated repression, we placed dCas9 expression under the optimized aTc-inducible promoter system (P*_tuf_*_(tetO-p)_ with P*_gap_*-*tetR*) and incorporated our RBS variants (G3C, G6T) to tune repression strength (Fig. 4e). Using NanoLuc as both target and readout, we observed distinct aTc-dependent response profiles consistent with their relative RBS strengths (Fig.4f). The stronger variants (native, G6T) exhibited switch-like repression (∼8-and ∼7-fold), whereas the weakest (G3C) generated a graded response with weaker repression (∼2-fold). Thus, these regulatory parts characterized above are portable to dCas9 and enable inducible and tunable CRISPRi. For subsequent experiments, we selected the G6T and G3C variants.

### CRISPRi-mediated inhibition of essential genes for biocontrol of engineered *B. longum* in tumors

For live bacterial therapeutics, control over the bacterial chassis itself could limit excessive proliferation or curtail bacterial activity once it is no longer needed^52^. Essential genes are required for bacterial growth and viability, but repressing them constitutively would impede bacterial propagation^53^. We therefore tested whether inducible CRISPRi-mediated repression of essential genes could inhibit *B. longum* growth on demand. We selected three genes representing distinct essential processes: *dnaA* (initiation of chromosome replication)^54^, *xfp* (fructose-6-phosphate phosphoketolase, a central enzyme of the bifid shunt)^25^ and *murC* (peptidoglycan biosynthesis)^55^, (Fig. 5a). Each was tested with two independent gRNAs and both dCas9 RBS variants, and their growth was monitored with and without aTc (G3C, Fig. 5b-e; G6T, Supplementary Fig. 6a-e). A non-targeting gRNA showed no differences in total growth, lag time, or growth rate upon aTc, confirming that neither aTc nor induced dCas9 affects growth. For targeting gRNAs, induction caused target-and guide-dependent growth inhibition, with major changes in total growth, lag time, and/or maximum growth rate. The translational context of dCas9 also affected growth inhibition. For example, strains with gRNA*_xfp_*_1_ produced a larger aTc-dependent growth inhibition when dCas9 carried G6T than G3C, again consistent with their relative RBS strengths.

**Fig. 5.**
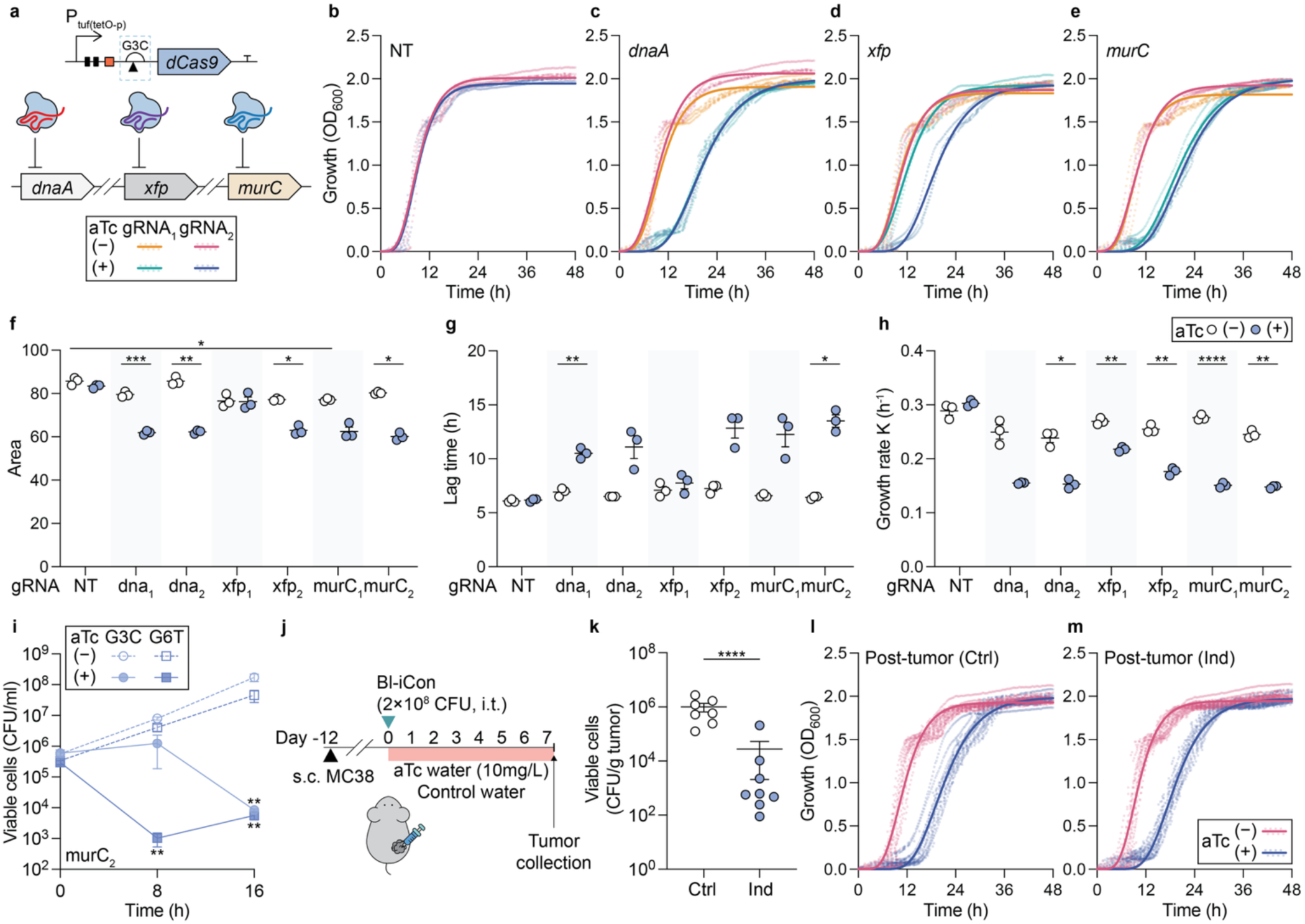
Inducible CRISPRi knockdown of essential genes controls *B. longum* growth in vitro and in tumors. (a–h) CRISPRi knockdown of essential genes with dCas9 expressed from the G3C RBS. See Supplementary Fig. 6 for the G6T RBS.| (a) Targeting of *dnaA* (BLON_RS00010), *xfp* (BLON_RS08935) and *murC* (BLON_RS04360), with two gRNAs each. (b–e) Growth curves in the presence or absence of 250 ng/ml aTc for a non-targeting gRNA (NT, b) or gRNAs targeting *dnaA* (c), *xfp* (d) or *murC* (e), fitted with the Gompertz growth model (n = 3). (f–h) Area under the growth curve up to 48 h (f), lag time (g) and Gompertz growth rate constant (K) (h) (n = 3). Brown–Forsythe and Welch ANOVA followed by Dunnett’s T3 multiple-comparison test on preselected pairs: each target versus NT at 0 and 250 ng/ml aTc, and 0 versus 250 ng/ml aTc within each target. (i) Viability of two *murC*-targeting strains (gRNA_murC2_) expressing dCas9 from two different RBSs (G3C and G6T, n = 3). Two-way ANOVA on log-transformed data followed by Tukey’s multiple-comparison test, comparing 0 vs. 250 ng/ml aTc within each strain at each time point. (j) Intratumoral injection of Bl-iCon (expressing gRNA_murC2_ and *dCas9* from the G3C RBS; 2 × 10^8^ CFU) into established subcutaneous MC38 tumors, with aTc-containing (Ind) or control (Ctrl) drinking water. Tumors were collected on day 7. (k) Bacterial burden in tumors (n = 8, pooled from two independent experiments). Welch’s t-test on log-transformed data. (l,m) Growth curves of colonies recovered from tumors and re-induced with aTc *in vitro* (n = 12). Colonies from control (Ctrl, l) and induced (Ind, m) tumors. See Supplementary Fig. 7 for growth parameters. Data are mean ± s.e.m. *P < 0.05, **P < 0.01, ***P < 0.001, ****P < 0.0001.

Among the targets and guides tested, gRNA*_murC_*_2_ produced robust growth inhibition upon aTc induction across both dCas9 variants, leading us to focus on this guide. Given the critical role of *murC* in peptidoglycan biosynthesis^55^, we sought to determine whether *murC* repression merely arrests growth or actively reduces bacterial viability. To distinguish growth arrest from killing, we quantified viable bacteria by CFU counting at 8 and 16h of aTc treatment (Fig. 5i). gRNA*_murC_*_2_-mediated repression caused killing rather than growth arrest, with the extent of viability loss depending on RBS strength. The stronger G6T RBS significantly reduced viability by ∼2.5 logs at 8 h, followed by partial regrowth at 16 h, whereas the weaker G3C RBS showed no significant change at 8 h and reduced viability only at 16 h. Considering durable suppression, we selected the G3C RBS with gRNA*_murC_*_2_ (hereafter the *B. longum* inducible containment, Bl-iCon) for *in vivo* evaluation to test whether inducible CRISPRi could enable externally triggered reduction of intratumoral bacteria.

We injected Bl-iCon intratumorally into established subcutaneous MC38 tumors and simultaneously provided aTc-containing or control drinking water (Fig. 5j). Seven days after injection, the aTc-treated group showed an ∼40-fold (∼1.6 log) reduction in intratumoral bacterial load relative to untreated controls (Fig. 5k). Bacteria recovered from the tumors of both aTc-treated and control mice remained responsive to aTc (Fig. 5i-m, Supplementary Fig. 7). Re-induction in culture significantly reduced total growth and growth rate while prolonging lag time. The magnitude of this response did not differ between the two post-tumor groups, and both exhibited the same aTc-dependent pattern as pre-tumor bacteria, indicating that the inducible growth-control phenotype remained stable seven days after tumor colonization.

Together, these results establish inducible CRISPRi as an externally controlled module for suppressing bacterial growth, and demonstrate that this control reduces bacterial burden within tumors—a key capability for biocontainment of engineered live bacterial therapeutics.

## Discussion

Effective use of live bacterial therapeutics for cancer requires not only tumor localization but also the ability to regulate bacterial activity after colonization^1–3^. *B. longum* is a tumor-homing probiotic with intrinsic antitumor activity^14–17,21^, yet its therapeutic potential has remained difficult to harness owing to the lack of established strategies for post-colonization regulation. Here, we address this limitation by bringing its gene expression and growth under external control. Building on a genetic foundation that supported stable plasmid maintenance and tunable gene expression (Fig. 1, Supplementary Fig. 1), we developed inducible modules for gene expression and CRISPRi-mediated repression of endogenous genes, including essential genes whose repression inhibited bacterial growth, and demonstrated that these modules remained functional not only in culture but also within tumors *in vivo* (Fig. 3, 5).

For an engineered bacterium to function *in vivo*, its genetic program should remain stable without selection, and its output should be reliably tunable to the desired level^22,23^. We achieved stable plasmid maintenance without continued selection using a native replicon identified from a human-derived *B. longum* isolate (Fig. 1a-b, Supplementary Fig. 1). We generated graded transcriptional and translational outputs by systematically substituting nucleotides within a native promoter and RBS, expanding the narrow range of constitutive expression previously available in this species^29^ (Fig. 1c-h, Supplementary Fig. 2). Notably, RBS variants retained their relative strengths when incorporated into an inducible system (Fig. 3f, 4f), suggesting that their graded behavior can extend across regulatory contexts. Together, these advances broaden the genetic tractability of *B. longum*.

Regulatory behavior established in culture cannot be assumed to persist after tumor colonization. The TME, characterized by hypoxia, nutrient limitation, immune pressure and spatial heterogeneity^56^, can alter bacterial physiology and, in turn, engineered regulatory systems^1,2,23^. We therefore evaluated our inducible systems in tumor-resident *B. longum*. In established tumors, heterologous gene expression (Fig. 3g-h) and CRISPRi-mediated growth control remained inducible (Fig. 5j-k). Growth control could be re-triggered in bacteria recovered from tumors following CRISPRi-mediated depletion (Fig. 5l-m, Supplementary Fig. 7), confirming that the system remained functional after *in vivo* passage.

The ability to reduce bacterial burden after administration is an important safety consideration for live bacterial therapeutics^22,52^. Inducible CRISPRi targeting the essential gene *murC* reduced viable bacterial burden in tumors by approximately ∼40-fold after induction, mirroring the progressive loss of viability observed in culture as cell-wall synthesis was suppressed (Fig. 5i-k). Because bacteria lacking an intact cell wall can increase bacterial susceptibility to host clearance^55,57^, *murC* repression may further promote immune-mediated elimination within the tumor, although we did not test this contribution directly. Thus, this system provides a potential route toward externally triggered biocontainment.

Separately, we identified signal peptides that supported extracellular release of payloads with prior therapeutic precedent in engineered bacteria, including SumIL2^17^, CCL21^7^, CXCL10^41^ and nanobodies^8^ targeting PD-L1 or CTLA-4 (Fig. 2g-h). Extracellular abundance varied with the signal peptide–payload pairing, highlighting the importance of matching secretion signals to individual cargos. Both wild-type *B. longum* and strains secreting either CCL21 or an anti-PD-L1 nanobody exhibited antitumor activity relative to PBS-treated controls (Fig. 2i-j). Although the engineered strains did not differ significantly from wild-type *B. longum* over the experimental period, except for a transient reduction in tumor volume in the Bl-CCL21 group at day 24, both engineered strains reached statistical significance relative to PBS-treated controls earlier than the wild type. These findings indicate that engineering the chassis for payload secretion did not abrogate its intrinsic antitumor activity. SumIL2, another payload secreted in this study, was previously shown to enhance antitumor efficacy beyond that of wild-type *B. longum*^17^. Taken together with this prior work, these findings support *B. longum* as a delivery chassis with intrinsic antitumor activity and suggest that additional therapeutic benefit may depend on selecting payloads that effectively augment this baseline activity.

Several limitations point to future directions. Although the plasmid system was stable under the conditions tested, chromosomal integration may prove preferable for translational use. Our *in vivo* experiments were limited to a single subcutaneous MC38 model, and evaluation in orthotopic settings and additional tumor types will be needed to establish generality. The long-term genetic and functional stability of the inducible and biocontainment systems, including performance during prolonged tumor residence or repeated induction, remains to be established. The growth-control system substantially reduced the intratumoral bacterial burden over the seven-day induction period but did not achieve complete elimination, leaving the factors that limit this control remain to be defined. Payload delivery and externally triggered containment were demonstrated in separate strains, and combining them will be an important next step toward more sophisticated living therapeutics.

Our study brings the gene expression and growth of *B. longum*, a gut symbiont with tumor-colonizing capacity, under external control after colonization. This control was not confined to tumors, as inducible expression was also retained in the gut (Fig. 3i-j, Supplementary Fig. 5c-j), the organism’s native niche, extending its utility beyond tumor-targeted applications. The functional CRISPRi established here opens the way to systematic perturbation of the genes underlying *B. longum* fitness and colonization in host-associated environments, as transposon screens in *B. breve*^46^ and CRISPRi screens *E. coli* in gut^58^ have shown. Targeted regulation of endogenous genes, including those governing metabolite production such as ILA (Fig. 4d), could enable causal studies of how *B. longum* physiology shapes host immunity. Together, these advances establish *B. longum* as a controllable therapeutic chassis and expand its utility for studying bacterial physiology in host-associated environments.

## Methods

### Bifidobacterium strains, culture conditions, and general procedures

*Bifidobacterium longum* ATCC 15697 (GenBank CP001095.1) was purchased from ATCC (Manassas, VA, USA). All other *Bifidobacterium* strains used to assess the host range of the native replicon pJL21, together with their sources and the antibiotic concentrations used for selection, are listed in Supplementary Table 1. Unless otherwise stated, *B. longum* ATCC 15697 was used throughout this study.

Wild-type *Bifidobacterium* strains were cultured in de Man, Rogosa, and Sharpe (MRS) medium (BD Difco) supplemented with 0.05% (w/v) cysteine-HCl and 100 mM potassium phosphate buffer (pH 7.3), hereafter referred to as modified MRS (mMRS). For engineered *B. longum* ATCC 15697 strains, mMRS containing 5 µg/mL chloramphenicol was used for routine cultivation. All *Bifidobacterium* strains were cultured at 37 °C under anaerobic conditions (5% H_2_, 5% CO_2_, 90% N_2_) in an anaerobic chamber (Coy Systems). Culture media and plates were pre-reduced overnight in the chamber before use.

Throughout this study, culture supernatants were clarified using a two-step centrifugation procedure, hereafter referred to as supernatant clarification. Cultures were centrifuged at 8,000 rpm for 10 min at 4 °C, after which the supernatants were transferred to fresh tubes and centrifuged again under the same conditions.

### Plasmid Construction

The native replicon pJL21 was amplified from *B. longum* MSK17.2 (obtained from the Symbiotic Bacterial Strain Bank, Duchossois Family Institute (DFI), University of Chicago) using primers jlD277 and jlD347. The sequence of pJL21, along with the sequences of all genetic parts, primers, and gBlocks used in this study, is provided in Supplementary table 4. All primers and gBlocks were synthesized by Integrated DNA Technologies (IDT) or Twist Bioscience.

Plasmids were constructed by Gibson assembly and restriction–ligation. Gibson assembly was performed using an in-house master mix prepared as previously described^59^. Restriction–ligation reactions were performed using restriction enzymes (BsaI and Esp3I, NEB) and T4 DNA ligase (NEB) according to the manufacturer’s instructions.

### Transformation

Assembled constructs were transformed into chemically competent *E. coli* S17 cells by heat shock at 42 °C for 1 min, followed by recovery in Luria–Bertani (LB) medium (BD Difco) at 37 °C for 1 h and plating on LB agar containing 25 µg/mL chloramphenicol. Plates were incubated overnight at 37 °C. Individual transformants were selected and cultured in LB medium supplemented with 25 µg/mL chloramphenicol at 37 °C with shaking at 250 rpm.

Bifidobacterial electrocompetent cells were prepared as previously described^17^. Briefly, 2 mL of an overnight culture was inoculated into 100 mL of mMRS containing 1% (w/v) glucose (1:50 dilution) and incubated at 37 °C for 6–10 h until an OD_600_ of 0.6–0.8 was reached. Cultures were chilled on ice and harvested by centrifugation at 8,000 rpm for 10 min at 4 °C. Cell pellets were washed twice with ice-cold wash buffer (0.5 M sucrose, 1.36 mM citric acid, pH 5.8), resuspended in 2 mL of storage buffer (wash buffer supplemented with 10% (v/v) glycerol), aliquoted, and stored at −80 °C until use. Assembled constructs or plasmids purified from *E. coli* were introduced into 50 µL of electrocompetent cells by electroporation at 25 µF, 200 Ω, and 2,000 V in a 0.2-cm-gap cuvette (Bio-Rad Gene Pulser Xcell). Electroporated cells were recovered in pre-reduced mMRS for 1 h at 37 °C under anaerobic conditions, then plated on RCM agar containing chloramphenicol (5 µg/mL for *B. longum* ATCC 15697; see Supplementary table 1 for other strains) and incubated anaerobically at 37 °C.

Individual transformant colonies were screened by colony PCR or plasmid extraction, and constructs were verified by Sanger sequencing (University of Chicago DNA Sequencing Facility) or whole-plasmid sequencing (GENEWIZ or Plasmidsaurus).

### Luminescence assay for *in vitro* cultures

Unless otherwise stated, overnight cultures were inoculated into fresh medium at a 1:10 dilution and incubated anaerobically at 37 °C for 8 h. Samples were diluted as needed, up to 10,000-fold, depending on the level of reporter expression. A 15 µL aliquot of each sample was mixed with 15 µL of Nano-Glo working reagent (Nano-Glo Substrate diluted 1:50 in Nano-Glo Buffer; Promega). Luminescence was measured using a microplate reader (Infinite 200 PRO, Tecan), using white flat-bottom 96-well plates (Corning) with an integration time of 1000 ms. OD_600_ was measured using 300 µL of the corresponding undiluted whole culture or post-treatment cell suspension before dilution. Luminescence was normalized to the corresponding OD_600_ value (RLU/OD_600_).

For evaluation of the aTc-inducible luciferase expression and CRISPRi-mediated repression systems, aTc was added at the indicated concentrations at the time of inoculation. For luminescence measurements of culture supernatants, samples were processed using the supernatant clarification procedure described above. For Proteinase K treatment, overnight cultures were washed twice with PBS and resuspended to the original culture volume in PBS. A 200 µL aliquot of the cell suspension was mixed with 800 µL of PBS. Samples were incubated at 37 °C for 4 h in the presence or absence of Proteinase K (Final concentration, 0.1 mg/mL; Qiagen).

### Plasmid copy number determination

Total DNA (genomic and plasmid) was extracted from overnight cultures using a microbial DNA extraction kit (DNeasy PowerLyzer Microbial Kit, Qiagen). Quantitative PCR was performed (QuantStudio 3, Applied Biosystems using SYBR Green master mix (Applied Biosystems) with primers targeting the chromosomal housekeeping gene *gyrB* (jlD949, jlD950) and the plasmid-encoded NanoLuc gene (jlD1265, jlD1266) (Supplementary table 4). Cycling conditions were 95 °C for 10 min, followed by 40 cycles of 95 °C for 15 s and 60 °C for 1 min. Melt-curve analysis was performed after amplification to confirm product specificity. Plasmid copy number was calculated using the ΔCt method and reported as plasmid copies per chromosome equivalent.

### Plasmid retention assay

Overnight cultures were inoculated into fresh medium at a 1:2,000 dilution in the presence or absence of 5 µg/mL chloramphenicol and grown to an OD_600_ of approximately 1.8 (∼11 generations per passage). Cultures were serially passaged under the same conditions for a total of seven passages. Luminescence and OD_600_ were measured at each passage before the subsequent subculture. At the end of the seventh passage, cultures grown in the absence of chloramphenicol were serially diluted and plated in parallel on RCM agar with and without 5 µg/mL chloramphenicol. Plates were incubated anaerobically at 37 °C for 2 days, and colony-forming units (CFU/mL) were determined.

### Flow cytometry

Overnight cultures were washed twice with PBS, resuspended in PBS, and incubated with aerobic shaking at 37 °C for 1 h to allow mCherry chromophore maturation. Cells were collected by centrifugation, fixed in 4% (w/v) paraformaldehyde in PBS for 10 min at room temperature, washed twice with PBS, and resuspended in PBS. mCherry fluorescence was measured using a BD LSRFortessa flow cytometer (excitation 561 nm; emission filter 610/20 nm), and at least 50,000 events were recorded per sample. Bacterial events were gated on FSC/SSC to exclude debris, and the mean mCherry fluorescence intensity of the gated population was used for analysis. Data were analyzed using FlowJo software (v10.8.1).

### Determination of transcription start sites by 5′ RACE

Bifidobacterial RNA was stabilized by mixing 500 µL of overnight culture with 1 mL of RNAprotect Bacteria Reagent (Qiagen), followed by incubation at room temperature for 5 min and centrifugation at 8,000 rpm for 10 min. Total RNA was extracted using the RNeasy Mini Kit (Qiagen) according to the manufacturer’s instructions, with a modified lysis buffer. Briefly, cell pellets were resuspended in 100 µL of lysis solution consisting of Proteinase K (2 mg/mL, Qiagen), lysozyme (20 mg/mL, Thermo Scientific), and mutanolysin (20 U/mL, Sigma) in TE buffer (10 mM Tris-HCl, 1 mM EDTA, pH 8.0), vortexed for 10 s, and incubated at 37 °C for 45 min. RNA was subsequently treated with TURBO DNase (Invitrogen) at 37 °C for 1 h according to the manufacturer’s protocol, purified using the RNA Clean & Concentrator-25 kit (Zymo Research), and eluted in 50 µL.

DNase-treated RNA was quantified by NanoDrop (Thermo Fisher), and up to 5 µg was used for gene-specific reverse transcription with SuperScript IV (SSIV; Invitrogen) and primer jlD1262 (specific to mCherry gene). Each reaction contained 10 µL of DNase-treated RNA, 1 µL of H_2_O, 1 µL of 10 mM dNTPs, and 1 µL of jlD1262 (20 µM). The mixture was incubated at 65 °C for 5 min and then placed on ice for 1 min. An SSIV master mix (4 µL of 5× buffer, 1 µL of 100 mM DTT, 1 µL of SSIV, and 1 µL of RiboLock RNase inhibitor (Thermo Fisher) per reaction was added, and samples were incubated at 55 °C for 90 min followed by inactivation at 80 °C for 15 min. 1 uL of RNase H was then added and samples were incubated at 37 °C for 30 min. Two reverse transcription reactions were performed per sample and pooled, purified using the DNA Clean & Concentrator kit (Zymo Research), and eluted in 20 µL.

For adapter ligation, 5 µL of cDNA and 2 µL of the single-stranded, 3′-blocked adapter jgD146 (40 µM) were incubated at 75 °C for 3 min and immediately placed on ice. A T4 RNA ligase master mix (2 µL of 10× buffer, 0.8 µL of DMSO or H_2_O, 0.2 µL of 100 mM ATP, 1.5 µL of T4 RNA ligase 1 (NEB), and 8.5 µL of 50% (w/v) PEG 8000 per reaction) was added, and reactions were incubated overnight at 22 °C. Two ligation reactions were performed per sample, pooled, and purified using the DNA Clean & Concentrator kit (Zymo Research). Ligated products were amplified by PCR using Q5 High-Fidelity DNA Polymerase (NEB) with primers jgD160 (forward) and jlD775 (reverse), purified using the DNA Clean & Concentrator kit and submitted for Sanger sequencing (University of Chicago DNA Sequencing Facility). Transcription start sites were determined by aligning the sequencing reads to the corresponding reference constructs and defining the 5′ end of the transcript as the first nucleotide immediately downstream of the ligated adapter sequence.

### Computational analysis

#### Substitution sensitivity analysis

Fluorescence of each single-nucleotide substitution variant was measured by flow cytometry. For each variant, fluorescence was normalized to that of the tuf wild-type sequence and expressed as a log_2_ fold change (log_2_FC). For each nucleotide position, the absolute log_2_FC values of all possible single-nucleotide substitutions at that position were averaged to yield the substitution sensitivity.

#### Identification of candidate promoters associated with highly expressed genes

Genes in *B. longum* NCC2705 were ranked according to their mean RPKM values across three RNA-seq samples (GEO accession GSE143410)^33^, and the 50 most highly expressed genes were selected. Putative operons were assigned manually based on the genome annotation of *B. longum* NCC2705 (GenBank AE014295.3). Consecutive genes on the same strand with short or overlapping intergenic regions were considered to belong to a single transcriptional unit. When multiple selected genes belonged to the same putative operon, only the promoter upstream of the first gene in the operon was retained, leaving 32 promoters.

For each of these genes, the entire intergenic region between its start codon and the 3′ end of the nearest upstream gene (70–510 bp) was taken as the promoter region. These regions were aligned to the *B. longum* subsp. infantis ATCC 15697 genome (GenBank CP001095.1) by BLASTn in Geneious Prime 2026, and orthology was confirmed by inspecting the gene immediately downstream of each aligned region. The aligned upstream regions in ATCC 15697 were designated candidate promoters associated with highly expressed genes. Four promoters were excluded, either because the NCC2705 promoter region could not be reliably aligned to the ATCC 15697 genome (n = 1) or because no corresponding gene was present downstream of the aligned region (n = 3), leaving 28 candidate promoters.

For five candidates whose promoters have been characterized previously (P*_tuf_*, P*_rplU_*, P*_rplM_*, P*_hup_*, and P*_gap_*)^60^, the reported −35 and −10 elements were used directly. For the remaining 23 candidates, the ATCC 15697 promoter regions were aligned to the *B. breve* UCC2003 genome (GenBank CP000303.1) by the same approach, and orthology was again confirmed from the gene immediately downstream of each aligned region. The −35 and −10 elements predicted previously for the corresponding UCC2003 promoters^61^ were then mapped onto the aligned ATCC 15697 sequences to assign the positions of the −35 and −10 elements. Four candidates were excluded because no −35 or −10 element had been predicted for the corresponding UCC2003 promoter, leaving 19 promoters assigned by alignment.

The −10 element was thus assigned for 24 promoters in total, comprising 5 previously characterized promoters and 19 assigned by alignment. For the −35 element, 5 of the 19 alignment-based promoters contained gaps or mismatches that precluded unambiguous assignment and were excluded from the −35 analysis. The −35 element was therefore assigned for 19 promoters in total, comprising 5 previously characterized promoters and 14 assigned by alignment. Sequence logos of the 6-nt −35 and −10 elements were generated separately using WebLogo3^62^.

#### Hybridization free energy calculation

The hybridization free energy between the 3′ end of the *B. longum* 16S rRNA (5′-CACCUCCUUUCU-3′) and the 25-nt region immediately upstream of the tuf gene start codon, containing the RBS and its single-nucleotide substitution variants (5′-CGAGACGUCC[AGGAGG]ACAAAAGUA-3′), was calculated using the DINAMelt Two-State Melting web server^63^. Each nucleotide within the bracketed 6-nt RBS was individually substituted with each of the other three nucleotides. Calculations were performed using the RNA (4.0) thermodynamic parameters at 37 °C in 1 M Na⁺ and 0 M Mg²⁺, with each strand at 10 µM. The predicted hybridization free energy (ΔG) was recorded in kcal/mol.

#### Signal peptide prediction

All annotated protein sequences of *B. longum* subsp. infantis ATCC 15697 (n = 2,551) were analyzed using SignalP 6.0^37^. Proteins with a predicted signal peptide probability of >0.99 were selected (n = 27), and their predicted signal peptide sequences were aligned using the Geneious alignment tool in Geneious Prime 2023. The consensus sequence derived from this alignment was used as a synthetic signal peptide (SP_syn_). In addition, four of the 27 predicted signal peptides (SP_2330_, SP_2335_, SP_5240_, and SP_8575_ from BLON_RS02330, BLON_RS02335 BLON_RS05240, BLON_RS08575, respectively) were used without modification. All signal peptide sequences are listed in Supplementary Fig. 3 and table 4.

#### Cell-wall anchoring module design

Cell-wall anchoring modules were derived from two LPXTG-containing surface proteins of *B. longum* ATCC 15697, encoded by BLON_RS01475 (655 residues) and BLON_RS11175 (1,179 residues). LPXTG motifs and the adjacent upstream disordered regions were identified from the UniProt annotations of the two proteins^39^. For each protein, three C-terminal modules of increasing length were tested: the region containing the LPXTG motif alone (residues 617–655 and 1146–1179, respectively), the same region extended to include the upstream disordered region (residues 587–655 and 1106–1179), and a further extended region (residues 491–655 and 1019–1179). Each module was fused to the C-terminus of NanoLuc via a (G₄S)₂ linker, with SP_2330_ fused to the N-terminus to direct translocation across the cytoplasmic membrane. All module sequences are listed in Supplementary table 4.

### Enzyme-linked immunosorbent assay (ELISA)

#### Sample preparation

Overnight cultures were inoculated into fresh medium at a 1:10 dilution and incubated anaerobically at 37 °C for 5-8 h. OD_600_ was measured from 300 µL aliquots of each culture for normalization. Cultures were placed on ice, supplemented with protease inhibitor solution at a 1:10 dilution (Roche), and clarified as described above. Clarified supernatants were stored at −20 °C until analysis.

#### Cytokine and chemokine quantification

Concentrations in culture supernatants were quantified using commercial Human IL-2, Mouse CCL21, and Mouse CXCL10 ELISA kits (R&D Systems) according to the manufacturer’s instructions. Before analysis, supernatants were diluted up to 512-fold in Reagent Diluent (R&D Systems) to obtain measurements within the assay range. Concentrations were corrected for the corresponding dilution factors and normalized to the OD_600_ of the original cultures.

#### Indirect His-tag ELISA

Relative nanobody concentrations were assessed by an indirect His-tag ELISA. Briefly, 100 µL of clarified culture supernatant was added to high-binding 96-well plates (R&D Systems) and incubated overnight at 4 °C. Wells were blocked with 1% (w/v) BSA in PBS for 2 h at room temperature. An anti-His-tag primary antibody (HIS.H8, Invitrogen; diluted 1:2000 in 1% BSA in PBS) was added and incubated for 2 h at room temperature. An HRP-conjugated anti-mouse IgG secondary antibody (Cell Signaling Technology; diluted 1:2000 in 1% BSA in PBS) was then added and incubated for 2 h at room temperature. Wells were washed three times for 5 min each with wash buffer (R&D Systems) between all incubation steps. TMB substrate was added, followed by stop solution (R&D Systems), and absorbance was measured at 450 nm with wavelength correction at 540 or 570 nm using a microplate reader (Infinite 200 PRO, Tecan). Signals were reported as OD₄₅₀ normalized to the OD_600_ of the original culture.

### Mouse experiments

All animal experiments were conducted in accordance with the Guide for the Care and Use of Laboratory Animals and approved by the Institutional Animal Care and Use Committee at the University of Chicago (ACUP No. 72610). All mice were maintained under a 12-h light/dark cycle at 20–23 °C and 30–70% relative humidity with ad libitum access to food and water.

#### Sample preparation

*B. longum* cultures were washed three times with PBS and resuspended in PBS containing 10% (v/v) glycerol to an OD_600_ of 0.7, corresponding to approximately 5 × 10^8^ CFU/mL. The suspensions were either aliquoted at this concentration or further concentrated 20-or 100-fold to approximately 1 × 10¹⁰ CFU/mL or 5 × 10^10^ CFU/mL, respectively, before aliquoting. Aliquots were stored at −80 °C and used within 1 week.

#### In vivo tumor models

Female C57BL/6 mice were obtained from Envigo, housed under specific pathogen-free conditions, and used at 8–10 weeks of age. MC38 cells were a gift from Dr. Jeffrey Hubbell and were cultured in DMEM supplemented with 10% FBS and 100 U/mL penicillin–streptomycin at 37 °C in 5% CO₂. Cells were routinely tested for mycoplasma contamination before use. MC38 cells (5 × 10^5^ cells in 100 µL of PBS) were injected subcutaneously into the flank of each mouse. Tumor dimensions were measured every 3 days with digital calipers, and tumor volume was calculated as (length × width × height)/2 mm³.

For therapeutic efficacy studies, treatment was initiated when tumors reached approximately 100 mm^3^. A 200 µL aliquot of wild-type or engineered B. longum suspension adjusted to an OD_600_ of 0.7 (1 × 10^8^ CFU per mouse) was administered intravenously three times at intervals of 3–4 days. Control mice received 200 µL of PBS intravenously according to the same schedule.

For inducible gene expression and CRISPRi studies, treatment was initiated when tumors reached 150–300 mm^3^. A 20 µL aliquot of the 20-fold-concentrated engineered *B. longum* suspension (2 × 10^8^ CFU per mouse) was administered as a single intratumoral injection. For induction, mice were provided with filtered drinking water containing 10 mg/L aTc, which was protected from light. At the experimental endpoint (day 3 or 7), tumors were collected and weighed before processing. Tumors were mechanically dissociated in 1–2 mL of PBS by pressing the tissue through sterile 40-µm cell strainers (Fisher Scientific) with a syringe plunger, and the resulting homogenates were used for subsequent analyses.

#### In vivo gut induction models

Germ-free Swiss Webster mice (male, ≥20 weeks old) bred and housed in the University of Chicago Gnotobiotic Research Animal Facility (GRAF) were used. Mice were orally gavaged with a bacterial suspension containing BL-iNL (approximately 3.3 × 10^9^ CFU per mouse) together with a defined microbial consortium (DMC), the composition of which has been described previously^45^, on days −9 and −6. Fecal samples were collected on days −3, 0, 1, 4, and 7. On day 0, fecal samples were collected before the initiation of aTc treatment. The experimental group was then provided drinking water containing aTc (10 mg/L), whereas the control group received drinking water without aTc. These conditions were maintained through day 7. On day 7, mice were euthanized and colonic contents were collected. Fecal samples and colonic contents were weighed before processing and homogenized in 0.5–1 mL of PBS using a PowerLyzer 24 Homogenizer (Qiagen) at 2,000 rpm for 5 min. Homogenates were allowed to stand for 5 min to permit large particulate matter to settle, and the upper fraction was collected for subsequent analyses.

#### Bacteria burden and luminescence measurement

To quantify the burden of engineered *B. longum*, homogenates were serially diluted and plated on RCM agar containing 5 µg/mL chloramphenicol. Plates were incubated anaerobically at 37 °C for 2 days, and colony-forming units were determined and expressed per gram of sample (CFU/g). For luminescence measurements, a 15 µL aliquot of homogenate was mixed with 15 µL of Nano-Glo working reagent and luminescence was measured as described above. Luminescence was normalized to the corresponding sample mass (RLU/g) or viable bacterial count (RLU/CFU).

#### Isolation of bacteria recovered from tumors

Colonies were randomly selected from the chloramphenicol-containing RCM agar plates used for CFU enumeration of tumor homogenates (3 colonies per tumor from 4 mice). Selected colonies were re-streaked on RCM agar containing 5 µg/mL chloramphenicol to ensure clonal purity. Single colonies were then inoculated into mMRS broth containing 5 µg/mL chloramphenicol and cultured for downstream analyses, including growth-curve measurements.

### Indole-3-lactic acid quantification

Overnight cultures were inoculated into fresh medium at a 1:10 dilution and incubated anaerobically at 37 °C for 8 h. After OD_600_ was measured from 300 µL aliquots, supernatants were clarified as described above. ILA abundance was determined using a customized metabolomics assay by liquid chromatography-mass spectrometry (LC-MS) (DFI Microbiome Metabolomics Platform, University of Chicago) and reported as relative abundance normalized to the corresponding culture OD_600_.

### Bacterial growth curves and viability assay

Overnight cultures were inoculated into fresh medium at a 1:2,000 dilution in a final volume of 200 µL per well in U-bottom 96-well plates (Corning). Cultures were grown in the presence or absence of aTc, which was added at the time of inoculation to a final concentration of 250 ng/mL.

For growth curves, cultures were incubated anaerobically at 37 °C for 48 h with continuous gentle shaking, and OD_600_ was recorded every 15 min using a microplate reader (Tecan Sunrise) with Magellan software. Growth curves were analyzed by calculating the area under the curve (AUC) and fitting a Gompertz growth model in GraphPad Prism (v11.0.2). Maximum OD_600_ was defined as the mean of the final four OD_600_ measurements, and lag time was defined as the time required for OD_600_ to reach 10% of this value.

For the viability assay, cultures were incubated anaerobically at 37 °C. At 0, 8, and 16 h, cultures were serially diluted and plated on RCM agar containing 5 µg/mL chloramphenicol. Plates were incubated anaerobically at 37 °C for 2 days, and viable colonies were expressed as CFU/mL.

### Statistics and reproducibility

All *in vitro* experiments were performed with at least three biological replicates, each shown as an individual data point where applicable. The number of independent experiments, the number of mice per group, and the statistical test used for each experiment are specified in the corresponding figure legends. Data are presented as mean ± SEM. Statistical analyses were performed using GraphPad Prism (v11.0.2). The statistical tests used for each analysis are specified in the corresponding figure legends. In general, two-group comparisons were analyzed using a two-tailed Student’s t-test, and multiple-group comparisons were analyzed using one-way or two-way ANOVA followed by an appropriate multiple-comparisons test. Tumor growth curves were compared using two-way ANOVA followed by Tukey’s multiple-comparisons test. CFU/mL and luminescence/OD_600_ values were log_10_-transformed before analysis. P < 0.05 was considered statistically significant, and significance is denoted as n.s. (not significant), *P < 0.05, **P < 0.01, ***P < 0.001, and ****P < 0.0001.

## Data availability

All data supporting the findings of this study are available within the paper and its Supplementary Materials. Key plasmid constructs generated in this study will be deposited in Addgene upon publication.

## Acknowledgements

We thank the following people for their help with this work: the Duchossois Family Institute (DFI) at the University of Chicago for providing human *Bifidobacterium* isolate strains; Yessenia Sierra and Catherine Hoyle at the University of Chicago Gnotobiotic Research Animal Facility for their assistance with gnotobiotic animal work; Jessica Little at the DFI Microbiome Metabolomics (DFIMM) core, University of Chicago, for metabolomics measurements; Jeffrey Hubbell for providing the tumor cell line; Ella Rotman and Jay Fuerte-Stone for blinding the samples; and Rory McGann and David Villegas for preparing the defined microbial consortia stocks.

## Funding statement

This work was supported by the National Institutes of Health, including the National Cancer Institute (R01CA292860 to R.R.W. and M.M.), the National Institute of General Medical Sciences (R35GM147478 to M.M.), and the National Institute of Allergy and Infectious Diseases (postdoctoral training grant T32-AI153020 to J.L.).

## Author contributions

Conceptualization, J.L., M.M.

Methodology, J.L., M.M.

Validation, J.L.

Formal analysis, J.L.

Investigation, J.L., J.G.

Resources, M.M.

Data curation, J.L.

Writing – original draft, J.L.

Writing – review and editing, J.L., J.G., R.W.W., M.M.

Visualization, J.L.

Supervision, M.M.

Project administration, M.M.

Funding acquisition, R.W.W., M.M.

## Competing interests

J.L. and M.M. are inventors on a provisional patent application related to this work filed by the University of Chicago. R.R.W. has stock and other ownership interests with Boost Therapeutics, Immvira, Reflexion Pharmaceuticals, Coordination Pharmaceuticals, Magi Therapeutics, and Oncosenescence. R.R.W. has served in a consulting or advisory role for Aettis, AstraZeneca, Coordination Pharmaceuticals, Genus, Merck Serono, Nano proteagen, NKMax America, and Shuttle Pharmaceuticals. R.R.W. has received research grant funding from Varian and Regeneron. R.R.W. has received compensation, including travel, accommodations, or expense reimbursement, from AstraZeneca, Boehringer Ingelheim, and Merck Serono. The other authors declare that they have no competing interests.

## Supplementary Information

Supplementary Figures 1-7

Supplementary Tables 1-4

## Notes

### Competing Interest Statement

J.L., R.R.W., and M.M. are inventors on a provisional patent application related to this work filed by the University of Chicago. R.R.W. has stock and other ownership interests with Boost Therapeutics, Immvira, Reflexion Pharmaceuticals, Coordination Pharmaceuticals, Magi Therapeutics, and Oncosenescence. R.R.W. has served in a consulting or advisory role for Aettis, AstraZeneca, Coordination Pharmaceuticals, Genus, Merck Serono, Nano proteagen, NKMax America, and Shuttle Pharmaceuticals. R.R.W. has received research grant funding from Varian and Regeneron. R.R.W. has received compensation, including travel, accommodations, or expense reimbursement, from AstraZeneca, Boehringer Ingelheim, and Merck Serono. The other authors declare that they have no competing interests.

