## Supplemental Data for "Programmable genetic control of tumor-colonizing *Bifidobacterium longum* for intratumoral therapeutic delivery and biocontainment"

### Supplementary Figures

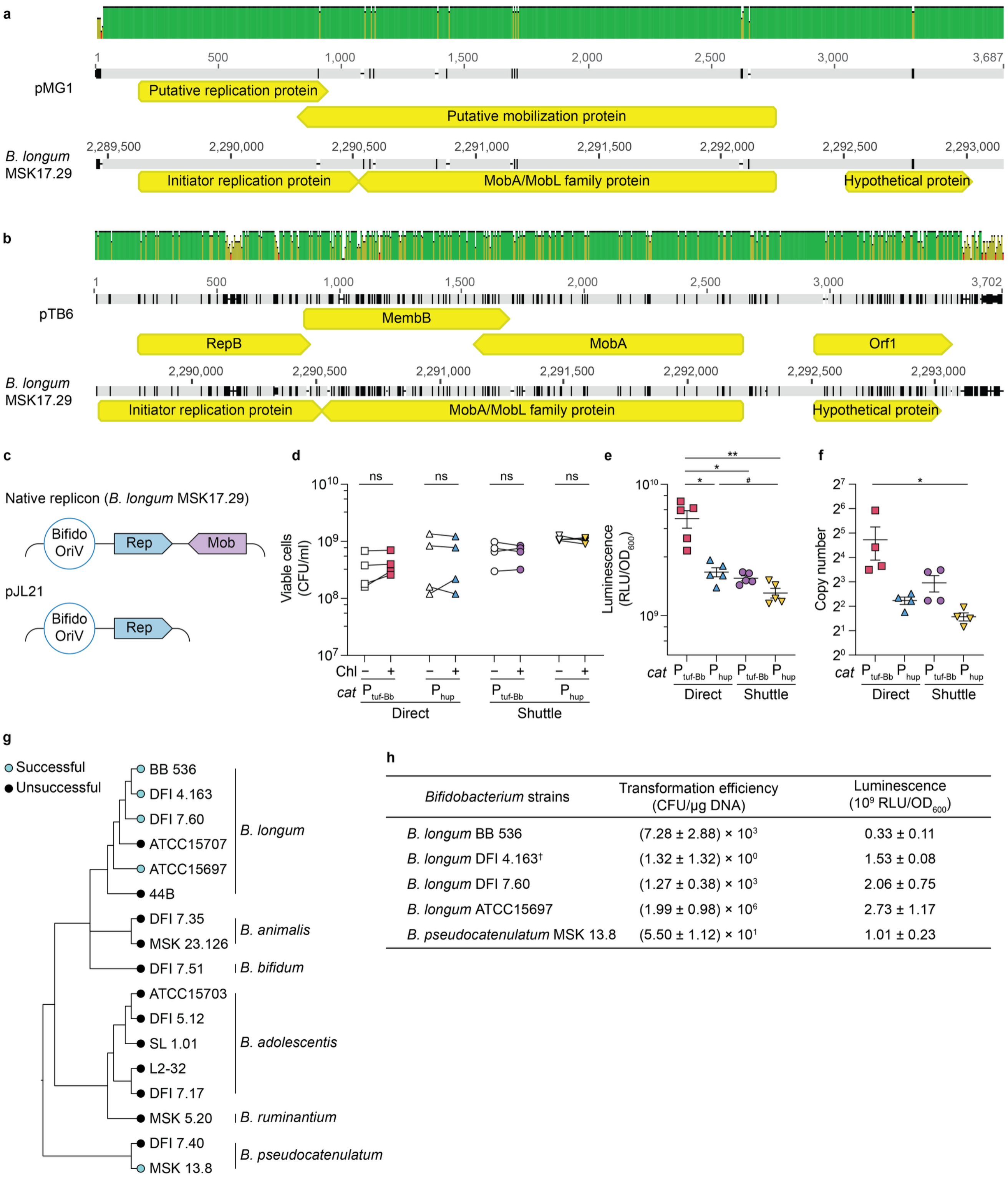


| *Bifidobacterium* strains | Transformation efficiency (CFU/μg DNA) | Luminescence ($10^9$ RLU/$OD_{600}$) |
|---|---|---|
| *B. longum* BB 536 | $(7.28 \pm 2.88) \times 10^3$ | $0.33 \pm 0.11$ |
| *B. longum* DFI 4.163† | $(1.32 \pm 1.32) \times 10^0$ | $1.53 \pm 0.08$ |
| *B. longum* DFI 7.60 | $(1.27 \pm 0.38) \times 10^3$ | $2.06 \pm 0.75$ |
| *B. longum* ATCC15697 | $(1.99 \pm 0.98) \times 10^6$ | $2.73 \pm 1.17$ |
| *B. pseudocatenulatum* MSK 13.8 | $(5.50 \pm 1.12) \times 10^1$ | $1.01 \pm 0.23$ |

**Supplementary Fig. 1. Native replicon-based plasmid system for *Bifidobacterium*.**

(a,b) Whole-sequence alignment of the native plasmids pMG1 (a) and pTB6 (b) against whole-genome sequencing data from *B. longum* MSK17.29.

(c) Schematic of the plasmid backbone pJL21. The native replicon identified in *B. longum* MSK17.29 (a,b) encodes an origin of replication (Bifido-OriV), a replication protein (Rep) and a mobilization gene (Mob). pJL21 comprises the Bifido-OriV and Rep elements of this replicon.

(d) Plasmid retention assay. After seven passages without chloramphenicol selection, cultures were plated on RCM agar with (Chl+) or without (Chl−) 5 µg/ml chloramphenicol, and CFU/ml was determined (n = 4). Two-tailed paired t-test.

(e,f) Luminescence per $OD_{600}$ (e) and plasmid copy number (f) in *B. longum* ATCC15697 for the indicated plasmid constructs (n = 5). Brown–Forsythe and Welch ANOVA on log-transformed data followed by Games–Howell's multiple-comparison test.

(g) Neighbor-joining phylogenetic tree of the *Bifidobacterium* strains tested with the native replicon-based plasmid system. The tree was inferred from *gyrB* nucleotide sequences aligned with MAFFT (v7.490) using Tamura–Nei distances and 500 bootstrap replicates. Branch lengths are not to scale. Strains are colored by transformation outcome (blue, successfully established with detectable luminescence; black, unsuccessful). Strains scored as unsuccessful were transformed at least three times with the Shuttle $P_{\mathit{tuf}\text{-Bb}}$*Cat* construct and additionally tested with the Direct $P_{\text{tuf-Bb}}$*Cat* construct.

(h) Transformation efficiency (CFU/µg) and luciferase expression (luminescence) for each successfully transformed strain. Unless otherwise indicated, values were determined using the Shuttle $P_{\mathit{tuf}\text{-Bb}}$*Cat* construct extracted from E. coli (n = 3). †, strains not transformable with the Shuttle $P_{\mathit{tuf}\text{-Bb}}$*Cat* construct, for which values were instead determined using the Direct $P_{\mathit{tuf}\text{-Bb}}$*Cat* construct extracted from *B. longum* ATCC15697 (n = 2).

Data in (e), (f) and (h) are mean ± s.e.m.; *P < 0.05, **P < 0.01, ***P < 0.001; #P = 0.053; n.s., not significant.

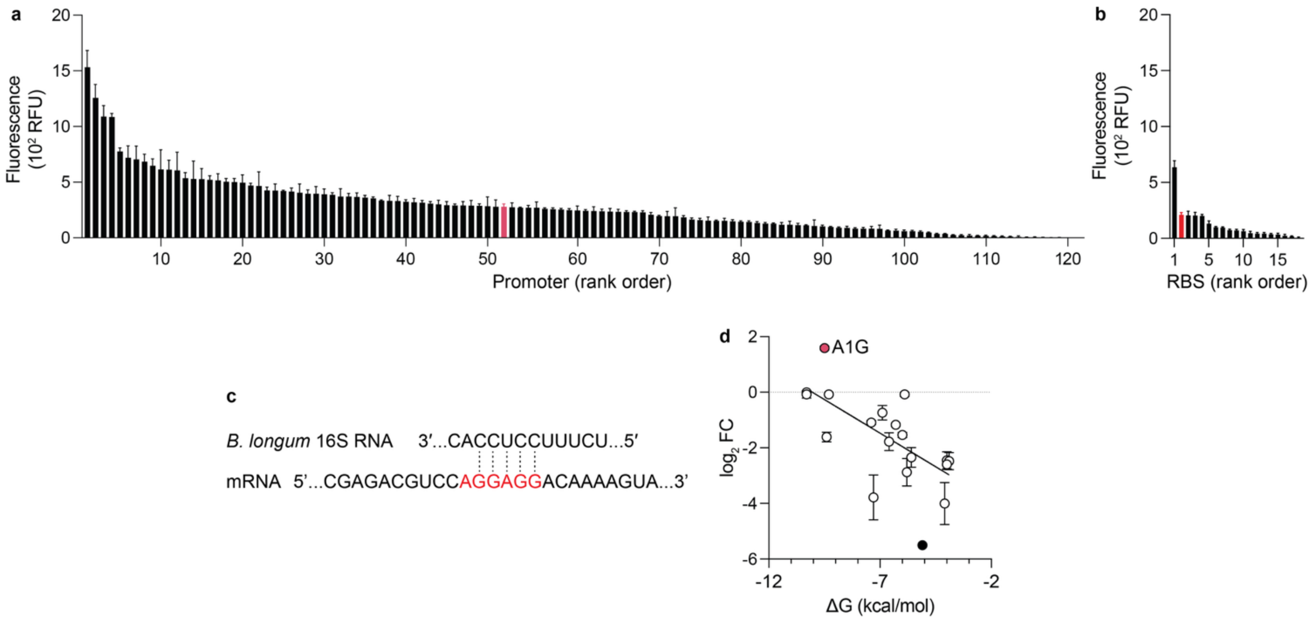


**Supplementary Fig. 2. Characterization of the synthetic promoter and RBS libraries in *B. longum*.**

(a,b) Fluorescence of the promoter (a, n = 3–4) and RBS (b, n = 3) libraries, shown as bar graphs in rank order.

(c) Base-pairing between the 3′ end of the *B. longum* 16S rRNA and the mRNA Shine−Dalgarno region used to calculate RBS−16S rRNA hybridization energy. Red nucleotides (AGGAGG) indicate the region varied in the RBS library.

(d) Relationship between expression ($\log_2$ fold change, $\log_2$FC) and the predicted RBS–16S rRNA hybridization energy. Expression was negatively correlated with hybridization energy (Spearman's $r = -0.72$, $p = 0.0004$). Red, the A1G variant, in which the first nucleotide was substituted from A to G; black, a variant with fluorescence below the detection limit, plotted at the floor value.

Data in (a), (b) and (d) are mean ± s.e.m.

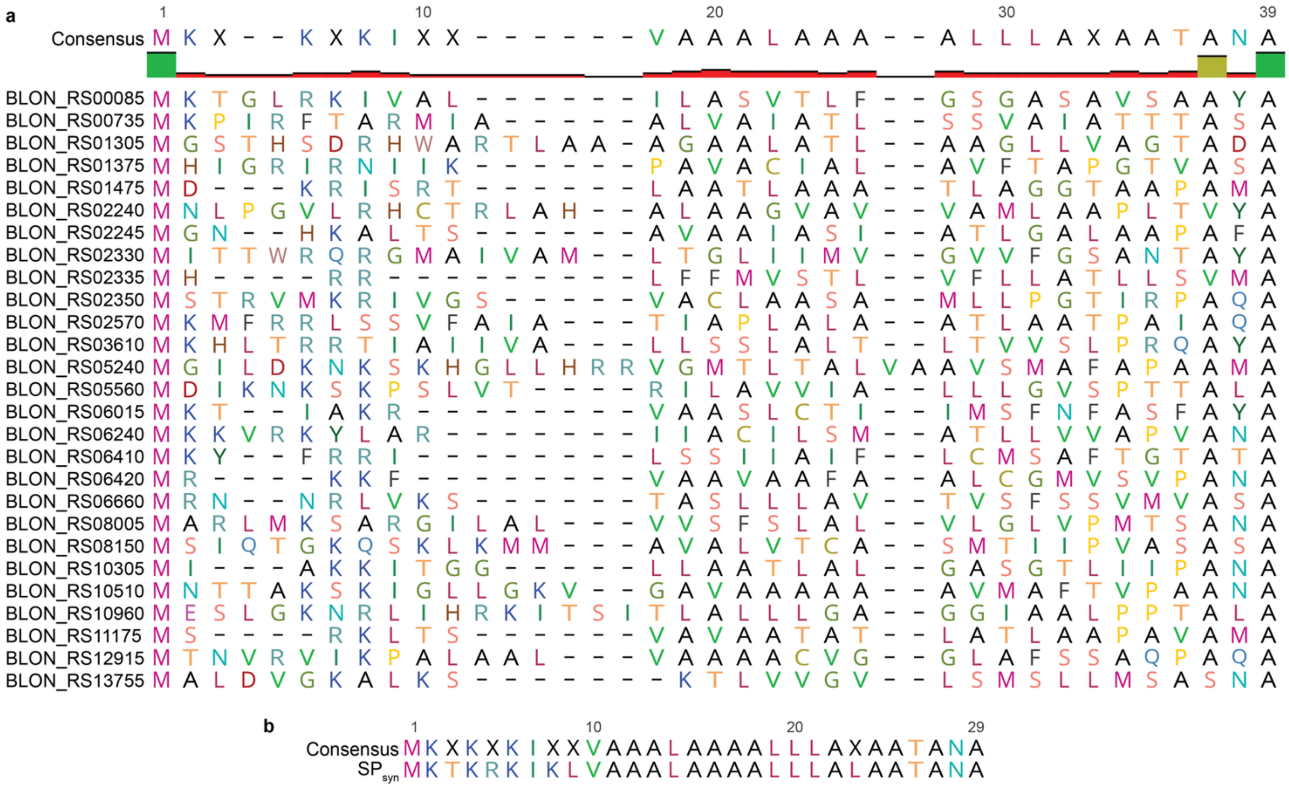


**Supplementary Fig. 3. Design of the synthetic signal peptide ($SP_{syn}$).**

(a) Alignment of 27 signal peptides scored >0.99 by SignalP 6.0.

(b) Consensus sequence of the alignment, used to define $SP_{syn}$.

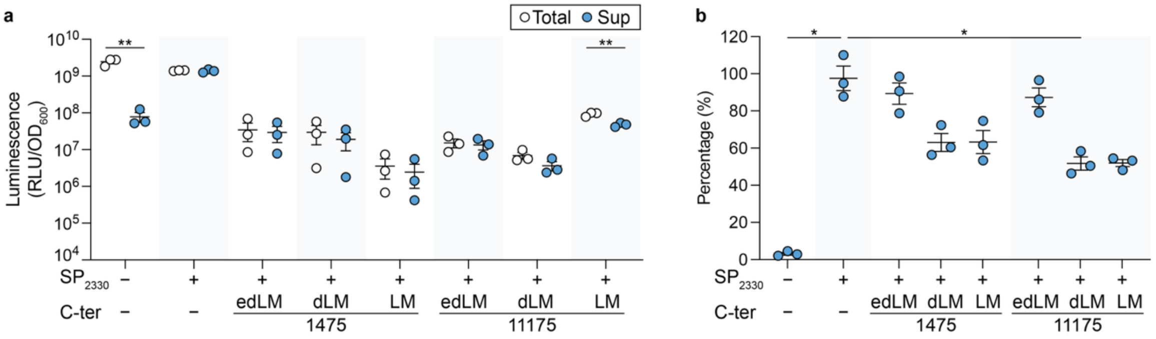


**Supplementary Fig. 4. Secretion of C-terminally anchored proteins into the supernatant.**

(a,b) Luminescence per $OD_{600}$ in whole culture (Total) and supernatant (Sup) for strains harboring the $SP_{2330}$ SP and a C-terminal LPXTG motif, $SP_{2330}$ alone (secretion control), and neither (cytosolic control) (a), and the percentage of secreted luminescence, calculated as (Sup/Total) × 100 (b) (n = 3). (a) Welch's t-test on log-transformed data. (b) Brown–Forsythe and Welch ANOVA followed by Dunnett's T3 multiple-comparison test against the secretion control.

Data in a and b are mean ± s.e.m., $^*P < 0.05$, $^{**}P < 0.01$.

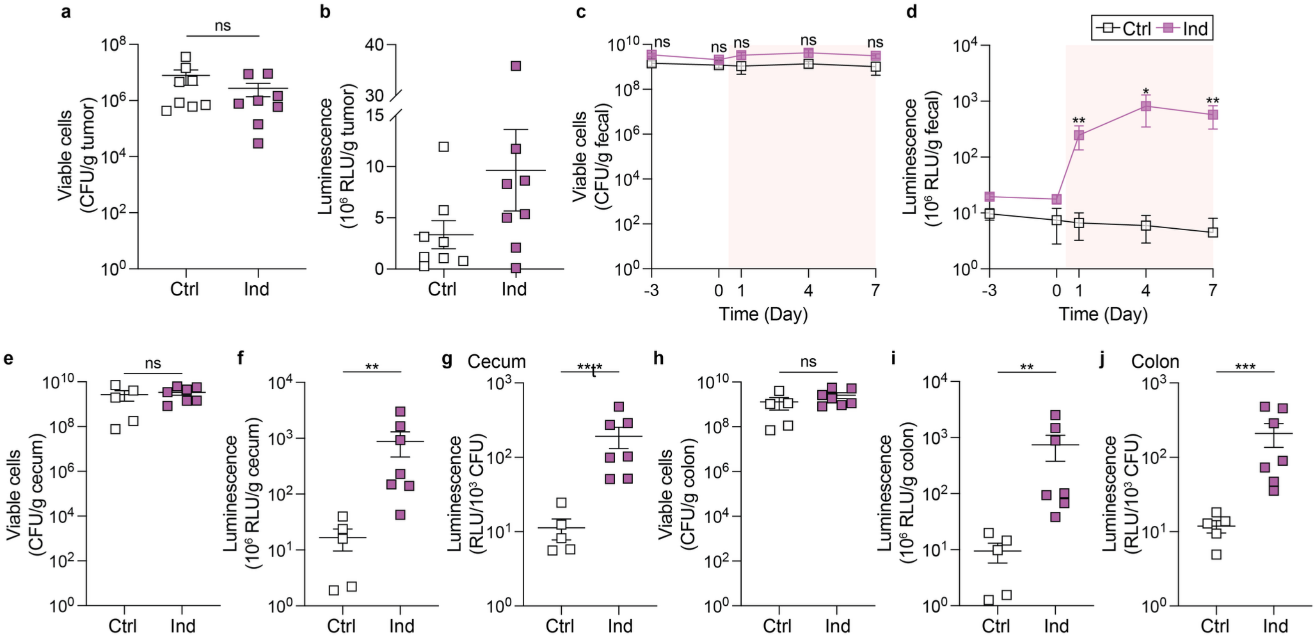


**Supplementary Fig. 5. Bacterial burden and luciferase activity of Bl-iNL in tumors and the gut.**

(a,b) Bacterial burden in tumors (a) and luminescence per g tumor (b) (n = 8, pooled from two independent experiments). (a) Welch's t-test on log-transformed data. (b) Welch's t-test.

(c,d) Bacterial burden in feces (c) and luminescence per g feces (d) over time (n = 5 for Ctrl and n = 7 for Ind, pooled from two independent experiments). Ctrl and Ind were compared at each time point by two-way repeated-measures ANOVA on log-transformed data followed by Šídák's multiple-comparison test.

(e–j) Bacterial burden and luminescence in the cecum (e–g) and colon (h–j): bacterial burden (e,h), luminescence per g (f,i) and luminescence per CFU (g,j) (n = 5 for Ctrl and n = 7 for Ind, pooled from two independent experiments). Welch's t-test on log-transformed data.

Data are mean ± s.e.m. *$P < 0.05$, **$P < 0.01$, ***$P < 0.001$, ****$P < 0.0001$; n.s., not significant.

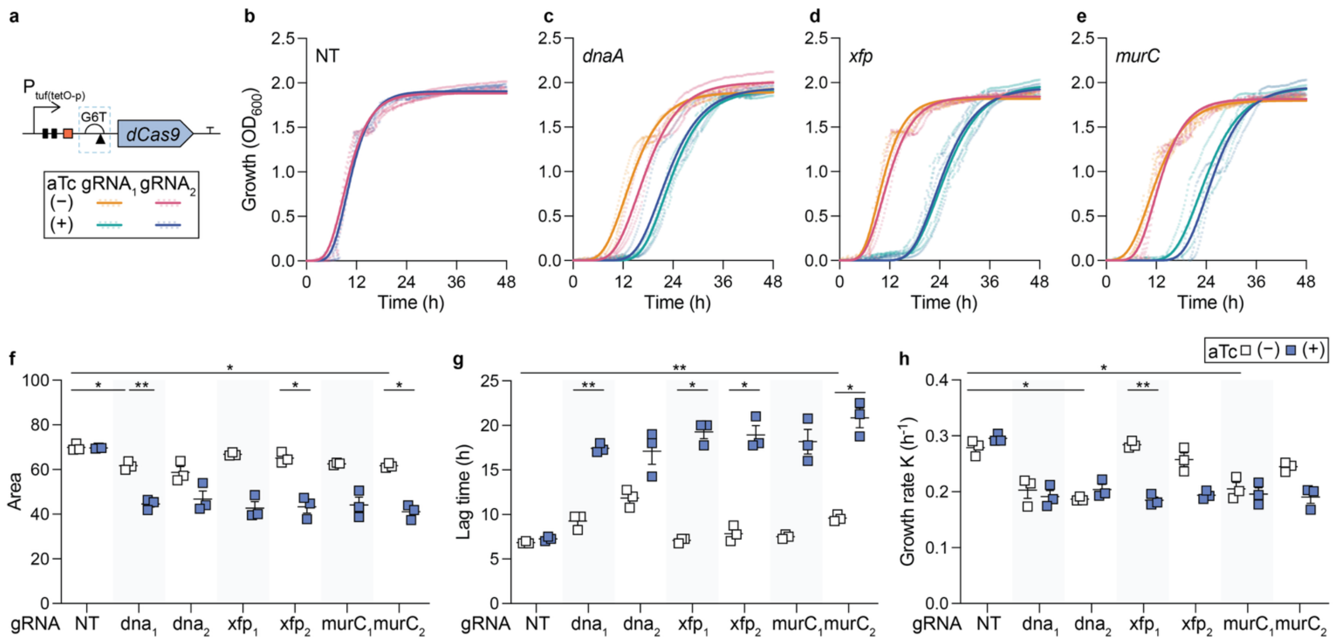


**Supplementary Fig. 6. CRISPRi knockdown of essential genes with dCas9 expressed from the G6T RBS.**

(a–h) As in Fig. 5a–h, but with dCas9 expressed from the G6T RBS. Targeting of *dnaA* (BLON_RS00010), *xfp* (BLON_RS08935) and *murC* (BLON_RS04360), with two gRNAs each (a).

(b–e) Growth curves in the presence or absence of 250 ng/ml aTc for a non-targeting gRNA (NT, b) or gRNAs targeting *dnaA* (c), *xfp* (d) or *murC* (e), fitted with the Gompertz growth model (n = 3).

(f–h) Area under the growth curve up to 48 h (f), lag time (g) and Gompertz growth rate constant (K) (h) (n = 3). Brown–Forsythe and Welch ANOVA followed by Dunnett's T3 multiple-comparison test on preselected pairs: each target versus NT at 0 and 250 ng/ml aTc, and 0 versus 250 ng/ml aTc within each target.

Data are mean ± s.e.m. *P < 0.05, **P < 0.01.

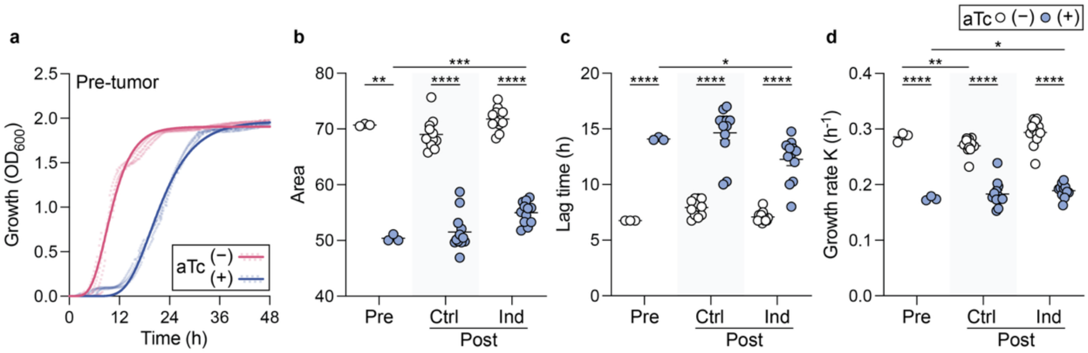


**Supplementary Fig. 7. CRISPRi-mediated growth attenuation is retained post-tumor.**

(a) Growth curves of the input (pre-tumor) Bl-iCon strain in the presence or absence of 250 ng/ml aTc.

(b–d) Area under the growth curve up to 48 h (b), lag time (c) and Gompertz growth rate constant (K) (d) for the pre-tumor strain (n = 3) and colonies recovered from control (Post, Ctrl, n = 12) or induced (Post, Ind, n = 12) tumors. Brown–Forsythe and Welch ANOVA followed by Dunnett's T3 multiple-comparison test on preselected pairs: each target versus NT at 0 and 250 ng/ml aTc, and 0 versus 250 ng/ml aTc within each target.

Data are mean ± s.e.m. *P < 0.05, **P < 0.01, ***P < 0.001, ****P < 0.0001.
